# Insights into G-Quadruplex–Hemin Dynamics Using Atomistic Simulations: Implications for Reactivity and Folding

**DOI:** 10.1101/2020.11.09.375691

**Authors:** Petr Stadlbauer, Barira Islam, Michal Otyepka, Jielin Chen, David Monchaud, Jun Zhou, Jean-Louis Mergny, Jiří Šponer

## Abstract

Guanine quadruplex nucleic acids (G4s) are involved in key biological processes such as replication or transcription. Beyond their biological relevance, G4s find applications as biotechnological tools since they readily bind hemin and enhance its peroxidase activity, creating a G4-DNAzyme. The biocatalytic properties of G4-DNAzymes have been thoroughly studied and used for biosensing purposes. Despite hundreds of applications and massive experimental efforts, the atomistic details of the reaction mechanism remain unclear. To help select between the different hypotheses currently under investigation, we use extended explicit-solvent molecular dynamics (MD) simulations to scrutinize the G4/hemin interaction. We find that besides the dominant conformation in which hemin is stacked atop the external G-quartets, hemin can also transiently bind to the loops and be brought to the external G-quartets through diverse delivery mechanisms. The simulations do not support the catalytic mechanism relying on a wobbling guanine. Similarly, catalytic role of the iron-bound water molecule is not in line with our results, however, given the simulation limitations, this observation should be considered with some caution. The simulations rather suggest tentative mechanisms in which the external G-quartet itself could be responsible for the unique H_2_O_2_-promoted biocatalytic properties of the G4/hemin complexes. Once stacked atop a terminal G-quartet, hemin rotates about its vertical axis while readily sampling shifted geometries where the iron transiently contacts oxygen atoms of the adjacent G-quartet. This dynamics is not apparent from the ensemble-averaged structure. We also visualize transient interactions between the stacked hemin and the G4 loops. Finally, we investigated interactions between hemin and on-pathway folding intermediates of the parallel-stranded G4 fold. The simulations suggest that hemin drives the folding of parallel-stranded G4s from slip-stranded intermediates, acting as a G4 chaperone. Limitations of the MD technique are briefly discussed.

**For Table of Contents Only:** 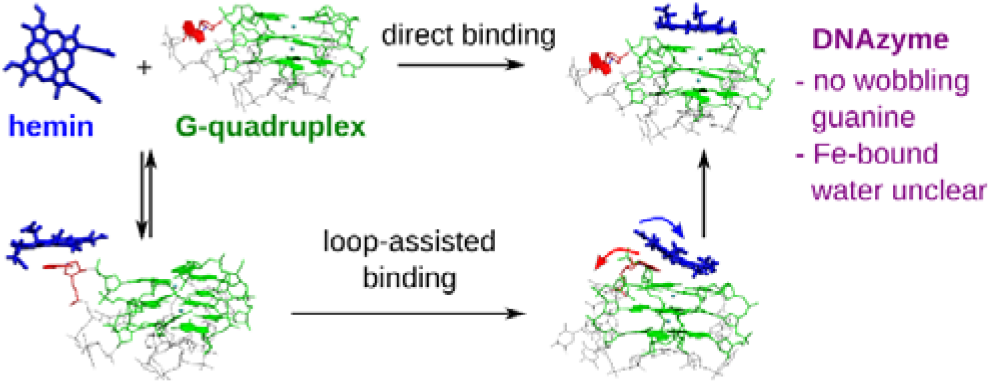

## INTRODUCTION

Guanine quadruplexes (G4s) are undoubtedly the most studied non-canonical nucleic acid structures. G4s are quadruple helices that fold from both DNA and RNA guanine (G)-rich sequences and adopt a variety of topologies depending on their nucleotide sequence and experimental conditions.^1–2^ The basic unit of G4 is a quartet of guanines (G-quartet), in which guanines are H-bonded in a cyclic arrangement (Figure 1A).^3^ G4s are created when at least two quartets stack upon each other to form a G-stem, which creates a channel running through its whole length, with inwardly pointing guanine O6 atoms that chelate cations (*e.g*., K^+^) and further stabilize the overall G4 architecture. G4 topology is characterized by structural rules comprising *syn*/*anti* conformations of guanines, strand directionality and loop types, which lead to a large topological diversity when combined.^4–6^

**Figure 1.**
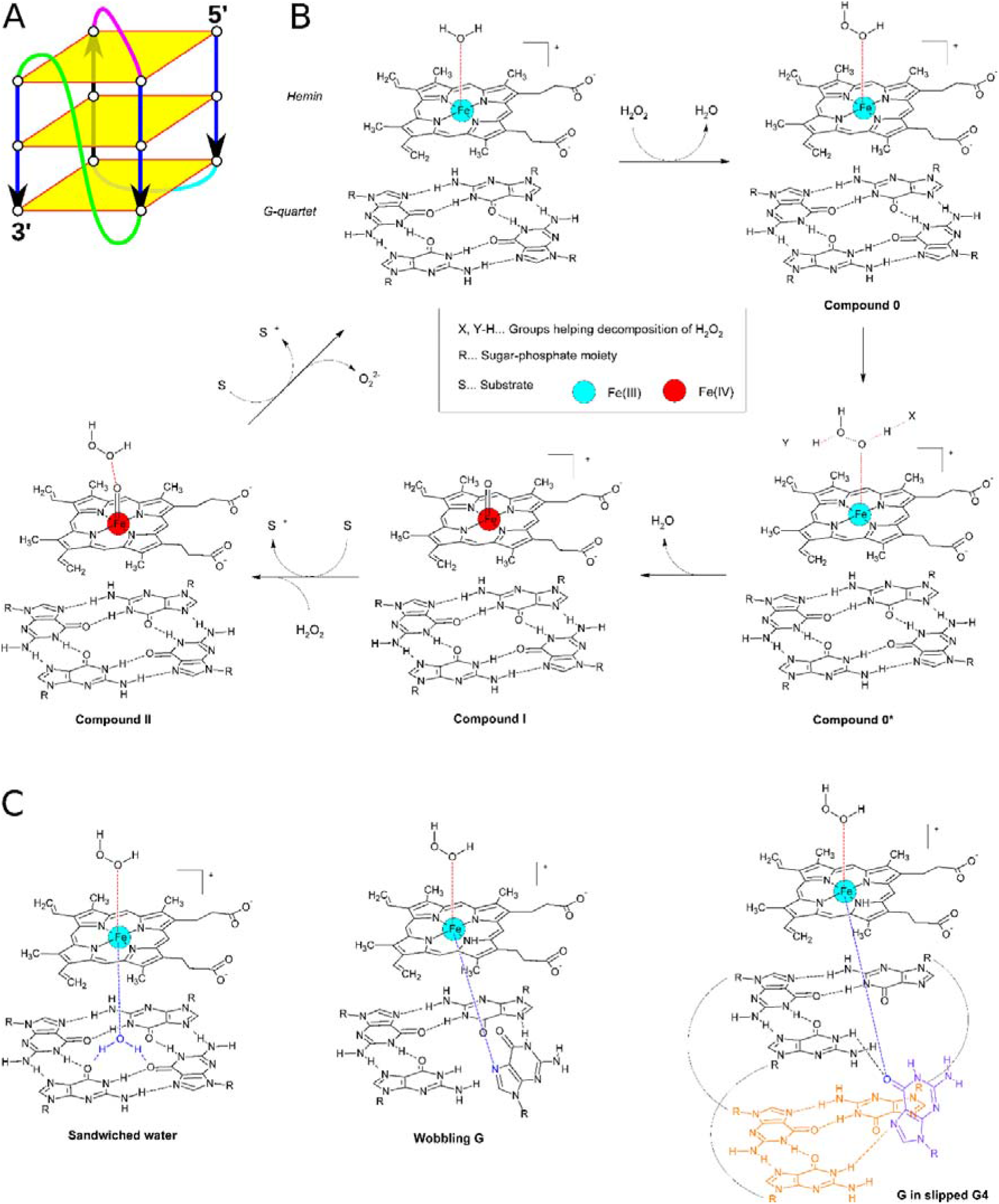
G4 structure, G4-hemin peroxidase cycle, and suggested hemin axial ligands. (A) G4 is formed by a G-stem, which is composed of stacked planar G-quartets (yellow; each composed of four Gs). Strands of the G-stem (blue) may be connected by loops: propeller (green), diagonal (purple) and lateral (cyan) are most common. (B) G4’s terminal G-quartet may bind hemin and form a DNAzyme with peroxidase activity. Its expected mechanism is thought to be similar to horseradish peroxidase and key steps are shown in the figure. (C) Axial ligands of iron in hemin bound to G4 suggested so far in the literature, shown at the Compound 0 state.^26–31^ In the rightmost structure, the nonplanar G (purple) is the first G of a downward-slipped strand, which tilts from its in-quartet position, formed with other three Gs (orange), upwards into the empty space available in the G-triad below the hemin.

Potential G4-forming sequences are widespread in the genome (>700,000 sequences),^7–8^ which lends credence to key G4 roles in cell’s life and functions.^1–2, 9^ This prevalence makes them active players in – and attractive targets for – genetic diseases including cancers^10–11^ and neuropathologies.^12–13^ G4s bind numerous ligands,^14^ including porphyrinic derivatives such as hemin (Fe(III)-protoporphyrin IX), which endows the resulting G4/hemin complex with unique catalytic properties referred to as G4-DNAzyme.^15^ The formation and H_2_O_2_-promoted peroxidase activity of G4-DNAzymes has found many applications notably as biosensors and for green chemistry purposes.^16–19^ G4-DNAzymes might also be relevant *in vivo* as heme is a naturally occurring cofactor involved in a variety of cellular processes. As intracellular heme levels vary in pathological conditions, G4-DNAzyme is hypothesized to cause oxidative damage to DNA.^20^ Alternatively, G4 might sequester free heme to protect cells against free heme-mediated toxicity.^21^

Despite dozens of applications now developed with G4-DNAzymes, structural information on the G4/hemin complexes is still fragmentary and the exact catalytic mechanism remains elusive. NMR experiments show the preferential stacking of hemin atop the 3′-quartet of parallel-stranded G4s.^22–24^ However, the dynamic aspect of G4-hemin binding has not been thoroughly addressed to date. The actual G4-DNAzyme mechanism is currently debated. Only recently, electron paramagnetic resonance spectroscopy (EPR) has indicated formation of Compound I (hemin cation-radical with oxygen bound to iron) as a reactive intermediate, which suggests a similarity to the horseradish peroxidase reaction mechanism (Figure 1B).^25^ Various hypothetical hemin activation mechanisms have been postulated, involving either a water molecule sandwiched in between the hemin and the adjacent quartet,^26–29^ the O6 atom of a wobbling G from the adjacent G-quartet,^30^ or G from a slipped G4 (Figure 1C).^31^ In the slipped G4 structure,^32–33^ the G could be tilted upwards towards hemin without causing any steric conflicts. All these catalytic intermediates rely on an oxygen or nitrogen atom as an activating axial ligand of the iron atom, but none of them has been unambiguously confirmed. In addition, reaction rate enhancements have been observed for G4 pre-catalysts with flanking bases^31, 34–37^ or when free nucleotides are added in the reaction solutions,^38–39^ possibly because they help with binding and activation of H_2_O_2_ in proximity of hemin iron (Figure 1B).

The G4-hemin interactions may also affect the G4 folding process. Recent studies suggest that G4s might follow intricate folding pathways best characterized by kinetic partitioning.^40–44^ Such a folding process is thought to be dominated by competition between diverse long-living G4 folds resulting from a combination of *syn*- and *anti*-oriented Gs.^41^ Many other structures can act as short-living transitory species during the folding process;^44–45^ in this line, MD simulations suggest an astonishingly large variety of structures that can participate in the G4 folding.^41^ The diversity of the structures and the expected large separation of time-scales between residency times of the metastable structures and transition times of active structural rearrangements make detection of transitory species by experimental techniques very challenging, opening space for advanced simulation studies.^44–45^ The G4 folding process may start with a very fast first phase resulting in an initial population of diverse G4 folds as long-living species.^44–45^ This population then equilibrates^40–41, 46–48^ over long time periods, often days,^40, 43, 49–50^ while some of the thermodynamically less stable but kinetically more accessible G4s act as long-living kinetic traps. Obviously, the kinetic accessibility of the different competing folds may depend on the initial ensemble of the molecules, while the lifetimes of the competing structures will depend on the temperature. Given the long time scale of the whole folding process, biologically relevant conformations of G4-forming sequences may differ from those in equilibrium.^41, 49^ In this context, ligands may substantially affect the complex G4-folding landscapes.^51–52^

Generally, transitions between different G4 topologies seem to require partial or complete unfolding of individual molecules. There is, however, one exception. Key folding intermediates of parallel-stranded all-*anti* G4s may correspond to transiently-populated slipped G4s with a reduced number of quartets.^32–33, 53^ Transitions between such structures can occur without any unfolding of the G-stem, through vertical strand-slipping movements. Although this process appears straightforward, experiments indicate that some slip-stranded structures can have long lifetimes for G4s with GGGG or longer G-tracts.^54^ Role of slip-stranded structures of parallel G4s has been considered in several recent experimental folding studies.^55–56^ As many ligands featuring a flat aromatic structure, including hemin, interact with (and seem to support) parallel-stranded G4s,^57–58^ they may affect the folding process, stabilizing some of the intermediates and/or modulating structural rearrangements.

In light of the unique catalytic properties of the G4/hemin complexes, we wonder whether the interactions of hemin with some of the folding intermediates could contribute to the G4-DNAzyme peroxidase activity.^31^ We thus used extensive all-atom explicit-solvent MD simulations to explore several aspects of the G4/hemin interaction. Our results show how hemin binds to parallel-stranded G4s and how the loops contribute to the binding process through an unprecedented delivery mechanism. We propose conformations that might contribute to the G4-DNAzyme catalytic activity and provide an explanation for the observed difference in activity of G4s with short loops, long loops and flanking nucleotides. Finally, we simulate partially folded slip-stranded parallel G4s complexed with hemin to investigate whether such intermediates can contribute to the overall catalytic activity and to understand how a ligand binding event may affect G4-folding pathway.

## METHODS

### Starting structures

We selected two intramolecular parallel-stranded all-*anti* G4s: the human telomeric G4 d[A(GGGTTA)3GGG] (PDB ID: 1KF1, X-ray structure; Figure 2),^59^ and d[TAGGGCGGGAGGGAGGGAA] (PDB ID: 2LEE, solution NMR structure, first frame).^60^ 1KF1 contains three three-nucleotide-long propeller loops and 2LEE contains three singlenucleotide-long propeller loops, and both G4s contain purine and pyrimidine bases in the loops. To assess how hemin interacts with bare G4 and possibly with its loops, we removed all the 5′- and 3′-flanking nucleotides. For binding simulations we placed a hemin molecule without bound axial ligands at a random position around the G4. We generated 20 starting positions for 1KF1 and 10 for 2LEE; the starting distance between G4 and hemin varied from contact distance to 40 Å with respect to the G4 center of mass (Supporting Figure S1).

**Figure 2.**
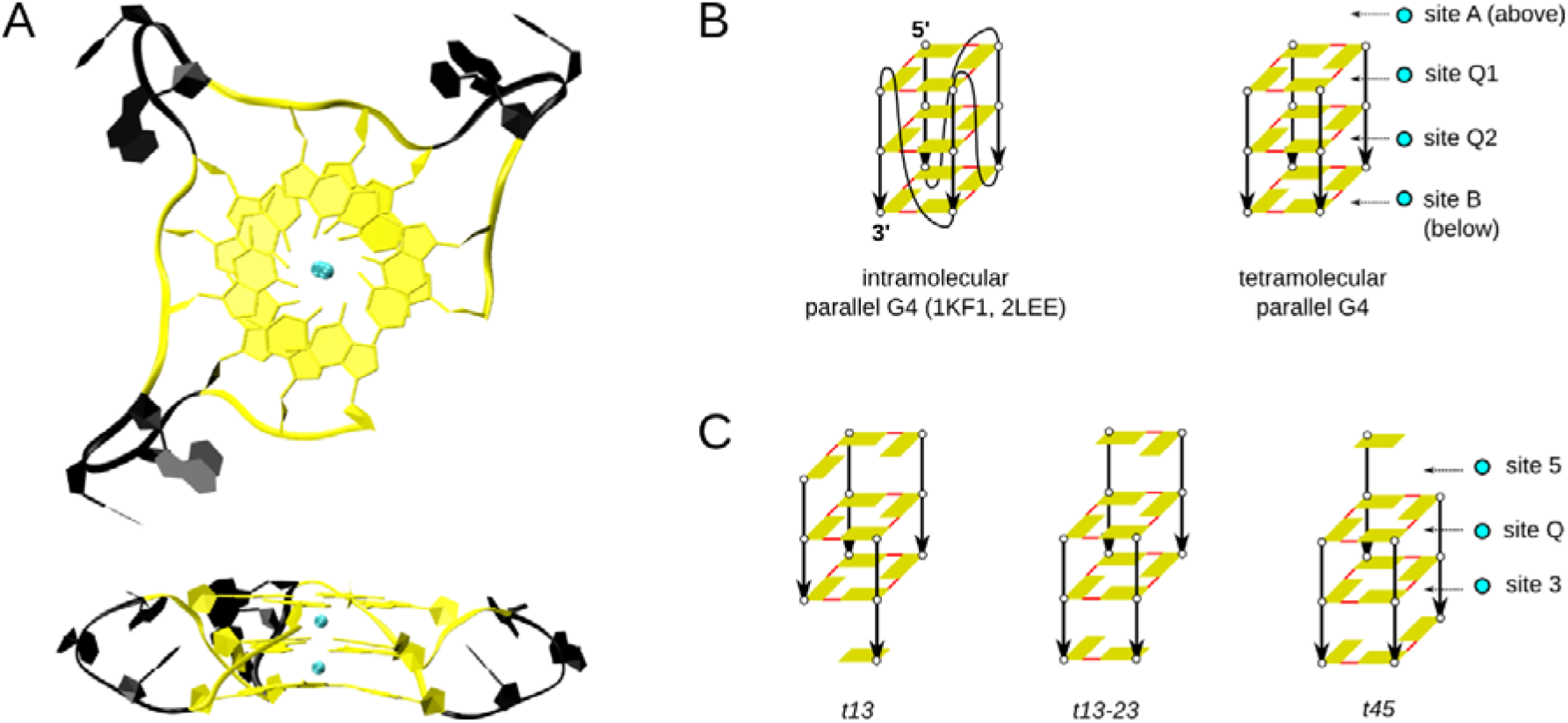
Starting structures in MD simulations. (A) top and side view of structure of the parallel-stranded G4 1KF1 (5′-flanking adenine present in the crystal structure was removed). The G-stem is depicted in yellow, TTA loops are shown in black, and potassium cations bound in the channel are cyan. (B) schemes of the intramolecular parallel-stranded G4s, and the tetramolecular parallel-stranded d[GGG]_4_ G4, shown along with designation of the channel binding sites. (C) d[GGG]_4_ G4s with one or more slipped strands, named as *t13*, *t13-23*, and *t45*, along with designation of the channel binding sites. Slip-stranded structures were suggested to act as late-stage folding/formation intermediates of parallel-stranded all-*anti* G4s.^32–33, 53^ The schemes in panels (B) and (C) and in next figures are drawn as follows: guanines are yellow; a pair of H-bonds between two guanines is depicted by a red line; the sugar-phosphate backbone is depicted by a black arrow showing the 5’→ 3’ progression; loop nucleotides and channel cations are not shown; bound hemin will be shown as a cyan rectangle (not present in this figure).

To assess whether a water molecule can act as a hemin-coordinating ligand and occupy a putative binding site between the hemin and the G4 channel, we took the simulated 1KF1 structure with hemin bound either atop the 5′ - or the 3′-quartet, removed the three TTA loops to create a three-quartet tetramolecular d[GGG]_4_ G4, and manually placed a water molecule in between the G4 and hemin. We also studied the hemin interaction with slip-stranded d[GGG]_4_ G4s: to this end, we took the d[GGG]_4_ G4 as described above and we manually prepared structures with slipped strands by removal and addition of Gs at appropriate ends, so that a G4 with two quartets was created. In doing so, we prepared three different structures (Figure 2), one with a strand slipped in the 3′-direction, designated *t13* (meaning tetramolecular G4, nominally slipped strand *1* in the 3′-direction), one with two neighboring slipped strands (*t13-23*, i.e. slipped strand *1* in the 3′-direction and strand *2* in the 3 ′-direction) and one with a strand slipped in the 5′-direction (*t45*; we use *t45* name instead of equivalent *t15* to better visually differentiate from *t13*). These three structures were simulated with and without bound hemin with initially three cations in the channel, at sites 5, Q, and 3 (Figure 2). Some simulations with 5′-bound hemin were initiated with only two channel cations, with the site 5 vacant.

We also investigated binding kinetics of hemin to G4s. For this task, we placed hemin farther away from G4s to avoid close initial contact of hemin with G4, to obtain more reliable results: i) we prepared three “large-box” systems with starting positions of hemin at a distance of about 80 Å from 1KF1, and ii) ten “midsize-box” systems with hemin placed at a distance of 30–45 Å from the three-quartet tetramolecular d[GGG]_4_ G4 (Figure S1).

### MD simulations

Each system was solvated in an octahedral box of SPC/E water molecules with the distance between solute and box border of at least 10 Å.^61^ When required by the hemin initial position, the box was enlarged to ensure that the hemin-G4 distance across the box border (to its periodic image) was at the desired length. Hemin and G4 concentration was about 0.01M in most simulations (except for the large-box and midsize-box binding simulations, see Supporting Tables S1 and S2). The simulation box was neutralized by K^+^ ions and then 0.15M KCl^62^ was added. Where intended, cations were placed inside the G4 channel. We applied the latest AMBER OL15 DNA force field.^63^ OL15 includes also several earlier modifications^64–66^ of the Cornell *et al*. force field.^67–68^ The overall combination of force-field parameters appears to be presently optimal for G4 simulations.^69^ GAFF with Shahrokh’s parameters was applied for hemin.^70^ The preparation was done in the leap module of AMBER16.^71^ Solvated structures were subjected to equilibration following standard protocol, described in Supporting Information. Production phase was performed in the CUDA version of pmemd module of AMBER16.^71–73^ SHAKE and SETTLE algorithms were used to constrain covalent bonds involving hydrogen.^74–75^ Hydrogen mass repartitioning scheme was used with integration time step set to 4 fs.^76^ Electrostatics was treated by the PME method and the cut-off for non-bonding interactions was set to 9 Å.^77–78^ Pressure was held at 1 bar and temperature at 300 K using the weak-coupling method.^79^ For the sake of completeness, in thirteen simulations dedicated to study kinetics of hemin-to-G4 association we used Langevin thermostat with a collision frequency of 2 ps^-1^ and Monte Carlo barostat. The reason for the change of thermostat is merely our decision to update our simulation protocol and these were the last simulations done for this study. We have recently compared both thermostats in G4 simulations and they appear to provide equivalent results.^80^

Length of individual simulations varied from 0.5 to 5 microseconds as specified in Table 1.

**Table 1.**
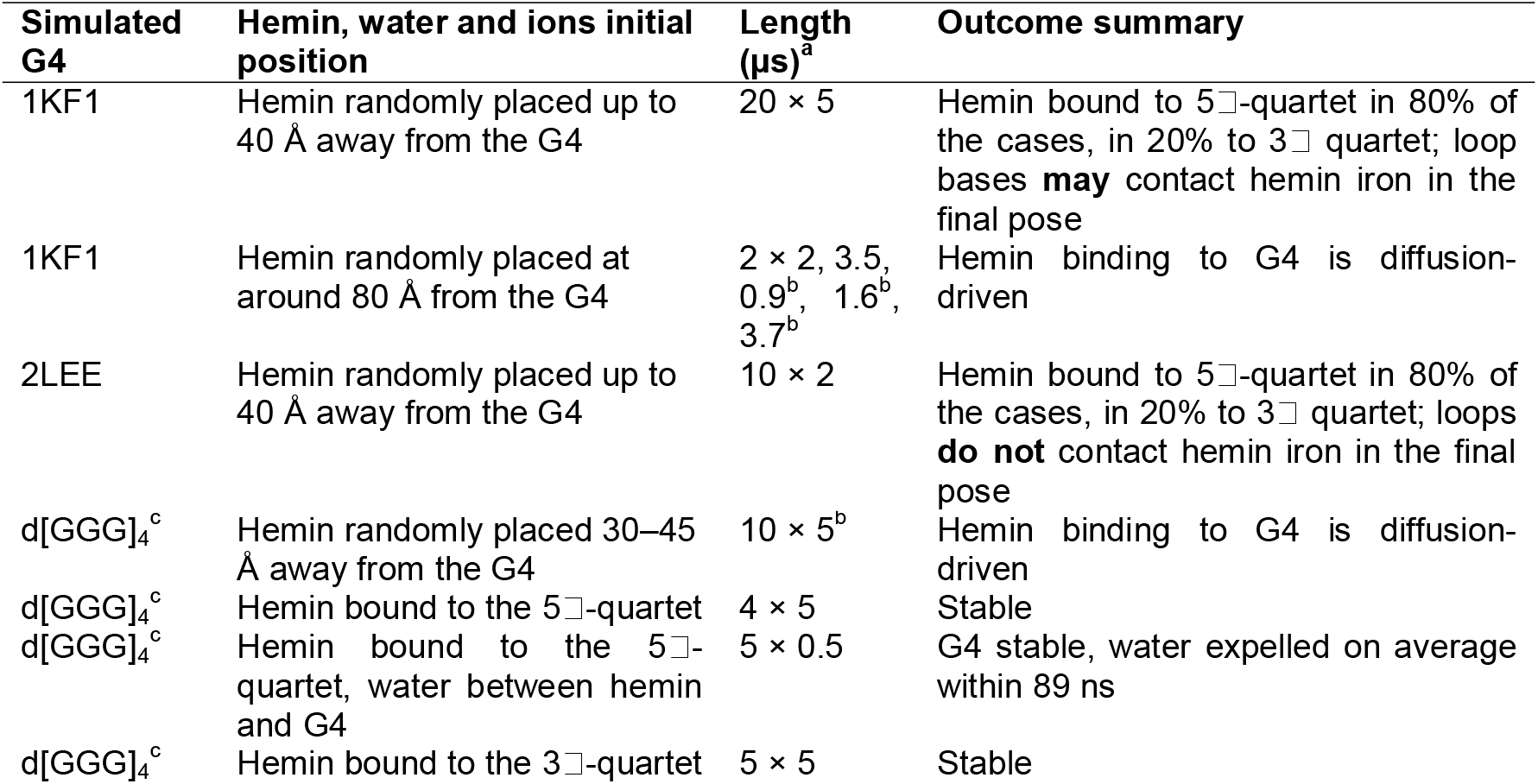
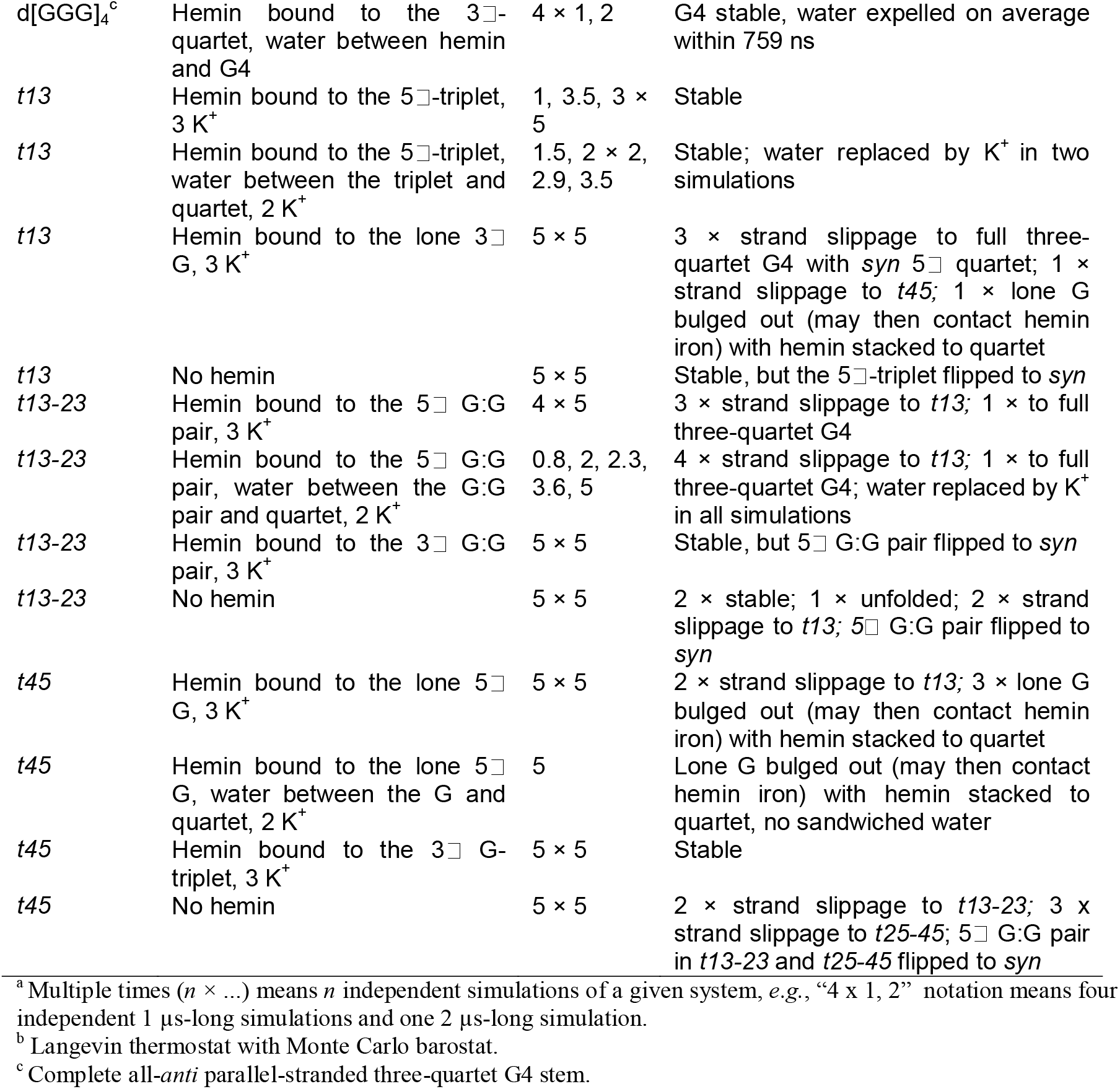
List of simulations.

| Simulated G4 | Hemin, water and ions initial position | Length (μs)<sup>a</sup> | Outcome summary |
| --- | --- | --- | --- |
| 1KF1 | Hemin randomly placed up to 40 Å away from the G4 | 20 × 5 | Hemin bound to 5□-quartet in 80% of the cases, in 20% to 3□ quartet; loop bases **may** contact hemin iron in the final pose |
| 1KF1 | Hemin randomly placed at around 80 Å from the G4 | 2 × 2, 3.5, 0.9<sup>b</sup>, 1.6<sup>b</sup>, 3.7<sup>b</sup> | Hemin binding to G4 is diffusion-driven |
| 2LEE | Hemin randomly placed up to 40 Å away from the G4 | 10 × 2 | Hemin bound to 5□-quartet in 80% of the cases, in 20% to 3□ quartet; loops **do not** contact hemin iron in the final pose |
| d[GGG]<sub>4</sub><sup>c</sup> | Hemin randomly placed 30–45 Å away from the G4 | 10 × 5<sup>b</sup> | Hemin binding to G4 is diffusion-driven |
| d[GGG]<sub>4</sub><sup>c</sup> | Hemin bound to the 5□-quartet | 4 × 5 | Stable |
| d[GGG]<sub>4</sub><sup>c</sup> | Hemin bound to the 5□-quartet, water between hemin and G4 | 5 × 0.5 | G4 stable, water expelled on average within 89 ns |
| d[GGG]<sub>4</sub><sup>c</sup> | Hemin bound to the 3□-quartet | 5 × 5 | Stable |
| d[GGG]<sub>4</sub><sup>c</sup> | Hemin bound to the 3'-quartet, water between hemin and G4 | 4 × 1, 2 | G4 stable, water expelled on average within 759 ns |
| *t13* | Hemin bound to the 5'-triplet, 3 K<sup>+</sup> | 1, 3.5, 3 × 5 | Stable |
| *t13* | Hemin bound to the 5'-triplet, water between the triplet and quartet, 2 K<sup>+</sup> | 1.5, 2 × 2, 2.9, 3.5 | Stable; water replaced by K<sup>+</sup> in two simulations |
| *t13* | Hemin bound to the lone 3' G, 3 K<sup>+</sup> | 5 × 5 | 3 × strand slippage to full three-quartet G4 with *syn* 5' quartet; 1 × strand slippage to *t45*; 1 × lone G bulged out (may then contact hemin iron) with hemin stacked to quartet |
| *t13* | No hemin | 5 × 5 | Stable, but the 5'-triplet flipped to *syn* |
| *t13-23* | Hemin bound to the 5' G:G pair, 3 K<sup>+</sup> | 4 × 5 | 3 × strand slippage to *t13*; 1 × to full three-quartet G4 |
| *t13-23* | Hemin bound to the 5' G:G pair, water between the G:G pair and quartet, 2 K<sup>+</sup> | 0.8, 2, 2.3, 3.6, 5 | 4 × strand slippage to *t13*; 1 × to full three-quartet G4; water replaced by K<sup>+</sup> in all simulations |
| *t13-23* | Hemin bound to the 3' G:G pair, 3 K<sup>+</sup> | 5 × 5 | Stable, but 5' G:G pair flipped to *syn* |
| *t13-23* | No hemin | 5 × 5 | 2 × stable; 1 × unfolded; 2 × strand slippage to *t13*; 5' G:G pair flipped to *syn* |
| *t45* | Hemin bound to the lone 5' G, 3 K<sup>+</sup> | 5 × 5 | 2 × strand slippage to *t13*; 3 × lone G bulged out (may then contact hemin iron) with hemin stacked to quartet |
| *t45* | Hemin bound to the lone 5' G, water between the G and quartet, 2 K<sup>+</sup> | 5 | Lone G bulged out (may then contact hemin iron) with hemin stacked to quartet, no sandwiched water |
| *t45* | Hemin bound to the 3' G-triplet, 3 K<sup>+</sup> | 5 × 5 | Stable |
| *t45* | No hemin | 5 × 5 | 2 × strand slippage to *t13-23*; 3 × strand slippage to *t25-45*; 5' G:G pair in *t13-23* and *t25-45* flipped to *syn* |

<sup>a</sup> Multiple times ( $n \times \dots$ ) means $n$ independent simulations of a given system, e.g., “4 × 1, 2” notation means four independent 1 μs-long simulations and one 2 μs-long simulation.
<sup>b</sup> Langevin thermostat with Monte Carlo barostat.
<sup>c</sup> Complete all-*anti* parallel-stranded three-quartet G4 stem.

## RESULTS

We have accumulated 483 μs of MD trajectories; their length and brief overall outcome are summarized in Table 1.

### Hemin binds to both 5′- and 3′-quartets

In the 20 simulations performed with 1KF1 and 10 simulations performed with 2LEE, hemin spontaneously bound to the two external G-quartets, according to three typical pathways: hemin could *i*-move directly to stack on the terminal quartets; *ii*-first make pre-binding to the loops and then be transferred to the stacked conformation by a conformational change of the loop; or *iii*-first interact transiently with the loops, be released back into the bulk solvent and then make another binding attempt. Regardless of the exact details of the binding process, in all simulations hemin eventually bound to the terminal quartets and stayed there until the end of simulation. For both 1KF1 and 2LEE, hemin bound to the 3□- and 5□-quartet in ~20% and ~80% of simulations, respectively (Figure 3A).

**Figure 3.**
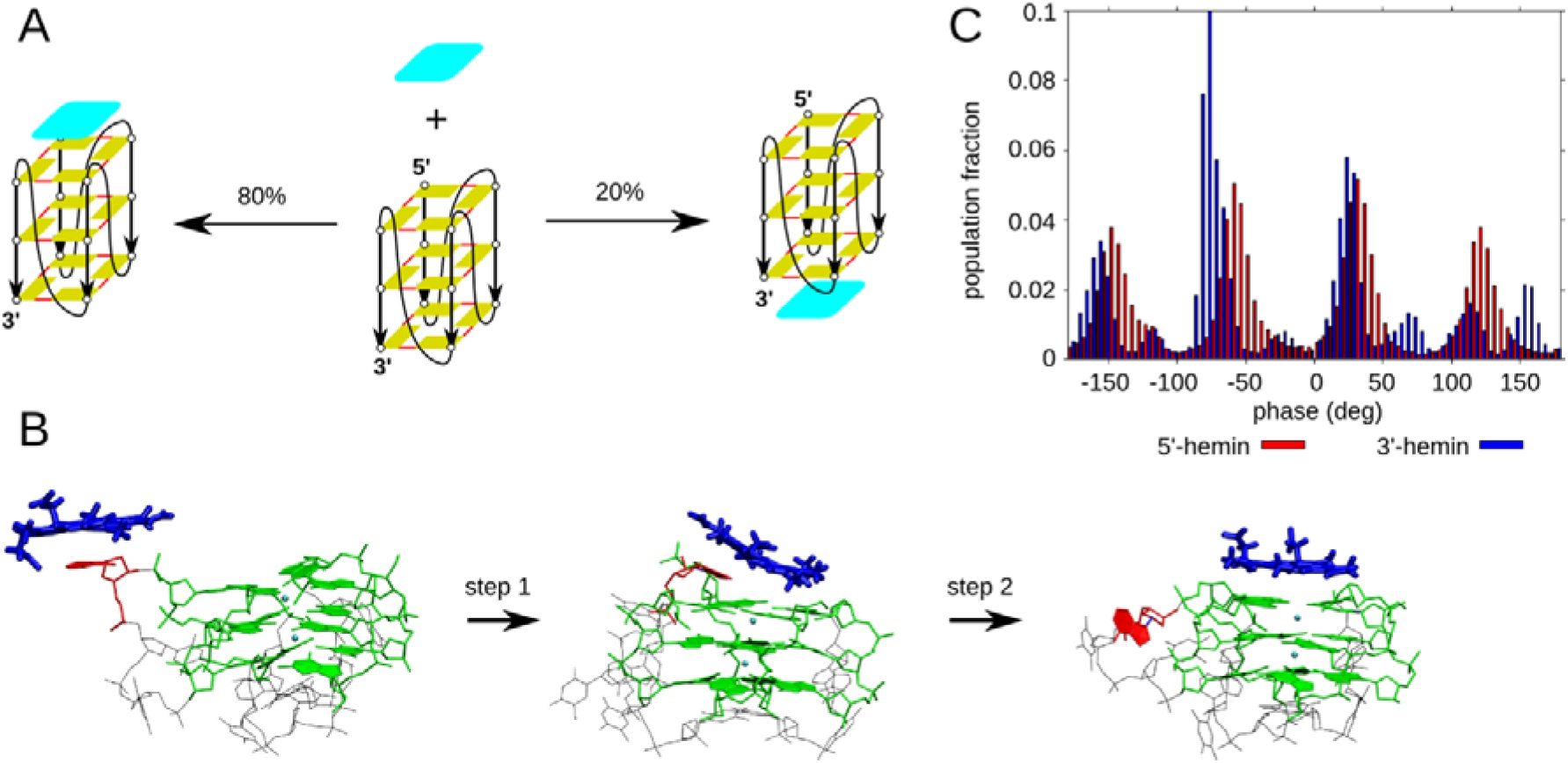
Binding of hemin to 1KF1 G4. (A) In all simulations hemin ended up bound either to the 5′- or 3′-terminal quartet, with a 4:1 probability ratio, similarly to the 2LEE simulations. The schematic representation is explained in the legend to Figure 2. (B) Adenine-assisted transfer of hemin to the 5′-quartet; hemin is shown in blue, the involved adenine in red, the G-stem is in green and other loop nucleotides appear in black. Once the first step happens, the second step is fast and takes only a few ns. The whole process is visualized in Supporting Movie SM1. (C) Rotational positioning of hemin stacked on terminal quartets is not uniform. Four-modal distribution was observed for hemin bound to 5′-quartet, and eight-modal to 3′ - quartet. Bin width was set to 5 deg. The phase is an arbitrarily chosen dihedral angle defined by the following four points: 1) Center of mass of C1□ atoms of G2 and G20 (*i.e*., guanines of the middle quartet from the first and last G-stretch), 2) center of mass of base heavy atoms of guanines forming the middle quartet, 3) hemin Fe(III), 4) hemin ring hydrogen atom located between the two carboxyethyl side chains.

Interception of the free hemin by 1KF1 G4, which contains three-nucleotide propeller loops, happened mostly *via* stacking to the loops. We observed four direct binding events to the 5′-quartet, three to the 3′-quartet and thirteen quartet stacking events mediated by the loops. Further, we detected nineteen transient bindings to the loops, after which the hemin departed back to the bulk. The three loops were involved in the binding approximately equally. As the three-nucleotide loops are flexible, we observed hemin stacking to a single base, formation of two-base stacking platforms, intercalation between two bases, as well as interaction with the grooves (Supporting Figure S2). Hemin usually remained bound to the loops for hundreds to thousands of ns, before it either unbound or slid to the 5′-quartet. A typical process looked like as if adenine (A) “picked hemin up” from bulk solvent and delivered it to the 5′-quartet (Figure 3B; Supporting Movie SM1): hemin first stacked on the loop’s A, then the loop underwent a conformational change and A with bound hemin moved above the 5′-quartet. Finally, hemin contacted the quartet and the A left the stacking position on the quartet, so that the hemin could stack atop the quartet through its entire surface (Figure 3B). No such hemin delivery process was found at the 3′-side, though we evidenced a single event in which hemin bound to a thymine (T) slid along the backbone to the 3′-quartet.

With 2LEE G4, hemin could bind directly to the terminal G-quartets or, more frequently, use pre-binding to the loops. We observed two direct bindings to the 5′-quartet, two to the 3′-quartet, six deliveries from the loops to the 5′-quartet and six transient bindings to the loops. Hemin bound only to either A8 or A12, but never to C4, probably due to its smaller stacking surface and also smaller sampling in the 2LEE simulation set (Supporting Figure S3). When bound to either of the As, hemin remained stacked usually for a few hundreds of ns, before it either unbound or slid from A to the 5′-quartet. Only direct binding events were found for the 3′-quartet.

### Hemin binding to G4 is diffusion-driven

To study hemin-G4 binding kinetics, we performed six large-box simulations of 1KF1 and ten midsize-box simulations of three-quartet tetramolecular G4 (d[GGG]_4_) with the initial distance between hemin and G4 of about 80 Å and 30–45 Å, respectively, to ensure no close contact of the two molecules. In case of 1KF1, we observed three direct bindings to the 5′-quartet, one direct binding to the 3′-quartet, and two loop binding events after which hemin was transferred to the 5′-quartet. Besides that, we observed two temporary binding events, one to a loop and one to the open groove lacking the loop. The picture is thus consistent with the smaller-box simulations. For d[GGG]_4_, we found five direct bindings to the 5′-quartet, two direct bindings to the 3′-quartet, three groove binding events after which hemin was quickly (a few ns) transferred to the 5′-quartet, and two temporary bindings to the backbone. Using the average time for binding to a quartet *t* and the concentration of hemin and G4 in a given simulation box *c*, using an approximate formula k = 1/*t*/*c*, the rate of binding would be estimated on the order of 10^9^ M^-1^ s^-1^, *i.e*., diffusion-driven, in both the large-box system (Supporting Table S1) and midsize-box system (Supporting Table S2). This suggests two conclusions: i) binding rate is not affected by simulation box size, provided it is big enough to avoid initial contact between hemin and G4, and ii) TTA loops of 1KF1 do not affect the overall rate of formation of G4-hemin complex in comparison to loop-free d[GGG]_4_; it means that while the loops of 1KF1 can bind hemin from bulk solvent faster than bare grooves of d[GGG]_4_, transfer of hemin from the loops to the terminal quartets is slower than transfer of hemin from the grooves to the terminal quartets by about the same factor.

### Dynamics of the bound hemin and its interactions with loop residues

Once stacked atop a terminal G-quartet, hemin rotated about its vertical axis. Since hemin has two sidechains and G4 has a sugar-phosphate backbone protruding from the plane of the terminal G-quartets, a four-modal distribution was found for hemin stacked on the 5′-quartet and eight-modal distribution for the 3′-quartet in 1KF1 (Figures 3C and Supporting Figure S4). The split from four-to eight-modal distribution was likely due to the interaction with the sugar rings exposed below the 3′-quartet. Hemin was not rigidly aligned exactly above the center of G4 stem, but rather wobbling around it due to thermal fluctuations (Figure 4A). The movement was more limited with hemin stacked atop the 3□-quartet. Adenines from TTA loops could rarely, and very briefly (at most a few ns), stack atop the hemin/G4 complex and form an A/hemin/5′-quartet sandwich. Interestingly, Ts located near the 3′-quartet could also form a similar sandwich (Figure 4B), whose formation was found more frequent than the 5′-quartet sandwich. The average frequency of formation was a few times per microsecond (counting only simulation time when hemin is stacked on the 3′-quartet). The T(O4)-iron direct contact time was up to dozens of ns, which would extend up to hundreds of ns if considering just the proximity of iron (6 Å). Therefore, the 1KF1/hemin complex allows for loop bases to appear in the vicinity of hemin central iron atom with a greater chance when hemin is stacked on the 3′-quartet.

**Figure 4.**
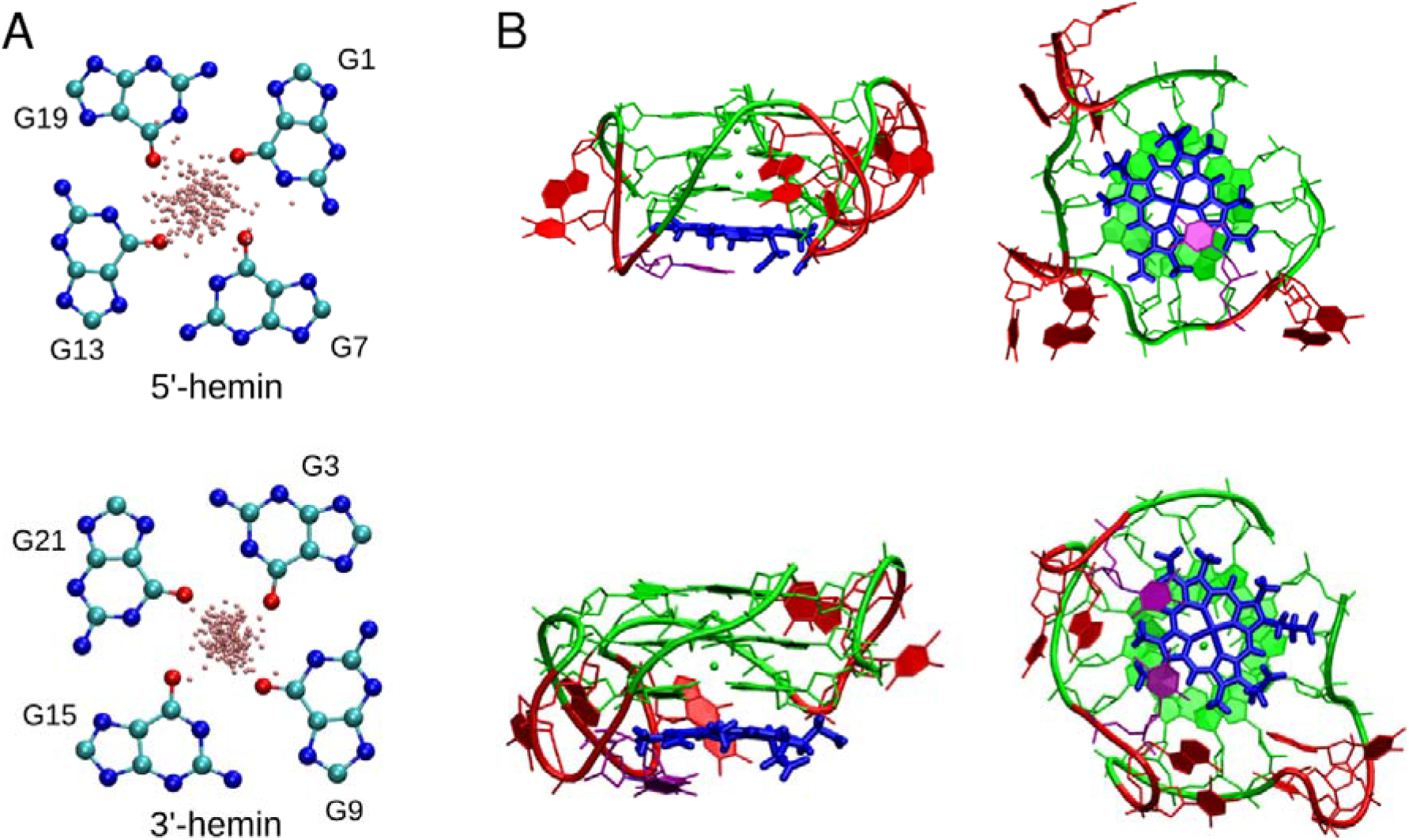
Hemin bound to the terminal quartets of 1KF1. (A) 200 randomly selected positions of the iron atom (pink) above the 5□- and 3□-quartet. (B) Iron in the center of hemin may occasionally be contacted by one of the loop thymines – structure with one thymine (top; two views) and structure with a T:T base pair (bottom; two views). Hemin is blue, the thymine(s) in the vicinity of iron purple, G-stem with channel cations green and other loop nucleotides red.

When bound to either of the 2LEE terminal G-quartets, hemin rotated with a four-modal distribution, caused by steric clashes between the hemin sidechains and the G4 sugarphosphate backbone (Supporting Figures S5 and S6). No 1KF1-like eight-modal distribution for 3′-binding was observed, because the short loops made the backbone shaped differently than in 1KF1. The wobbling movement of hemin around the G-stem axis was observed again. Due to steric constraints, none of the single-nucleotide loops could stack, even transiently, above or below the hemin to form a sandwich-like structure and reach the central iron atom of hemin. Therefore, the hemin rotation was not affected, and, more importantly, the loops’ bases are unlikely to contribute to the catalytic properties of the G4/hemin system, should the reactive conformation be the one with hemin fully stacked on G4.

No water molecule remained trapped between freshly stacked hemin and the terminal G-quartet in any of the binding simulations. All water molecules moving around were always expelled before hemin fully stacked atop the quartet. No ion exchanges of the two channel cations bound inside the G-stem of 1KF1 and 2LEE with the bulk were observed.

### A water molecule between the hemin and the G4 is expelled

Subsequently, we performed simulations of d[GGG]_4_ with hemin bound already in the initial structure. The G4 was entirely stable in all simulations while hemin was again rotating about its vertical axis (Supporting Figures S7, S8 and S9). When stacked to the 5′-quartet, the rotation had a four-modal distribution (Figures S7 and S9). Hemin was again wobbling around the G4 axis. Two K^+^ ions remained in the channel at sites Q1 and Q2 (see Figure 2) at all times; no ion exchanges were observed. With hemin stacked on the 3′-quartet, the rotation was characterized by a four-modal distribution (Figures S8 and S9), with four main peaks and four shoulders (Figure S9).

Since we did not observe any water molecule spontaneously occupying the space in between the hemin and the G4 in our simulations, we carried out ten simulations starting with a water molecule manually placed between the d[GGG]_4_ and either the 5′ - (water at site A) or the 3′-stacked hemin (site B). In all simulations performed with the 5′-stacked hemin, the water molecule quickly (within 1-286 ns) escaped into bulk, while the lifetimes were longer with the 3′-stacked hemin (217-1018 ns). The water molecule always left the G4 by transient incorporation into the terminal quartet *via* H-bonding,^80^ which was followed by a quick (order of ns) expulsion to the groove. Within the force-field approximation, the binding of a water molecule in between the hemin and the G4 is thus a very unlikely event, even if we do not have any estimate of its free-energy penalty (as no water binding event was observed). The different expulsion times at 5′ - and 3′ - ends may be explained by different G-quartet buckling: in both 1KF1 and the related d[GGG]_4_, both in the crystal structures and in the simulations, the 5′-quartet is nearly perfectly flat even without stacked hemin; the 3′-quartet without stacked hemin is buckled towards the G4 center (Supporting Figure S10), and hemin binding makes it flat. However, in the simulations with a water molecule initially placed at site B, the buckling re-occurs and the water molecule has thus a little more available space.

### Hemin stabilizes slipped G4s and may accelerate transition to the complete stem

Next, we simulated hemin bound to slip-stranded G4s, suggested late-stage on-pathway intermediates in folding of parallel-stranded G4s.^32–33, 41^ The goal was two-fold, *i*-to see if some slip-stranded structures can sample potential catalytically potent geometries (*i.e*., geometries where hemin may avoid perfect stacking to a quartet or geometries with base functional groups in proximity of the iron) and *ii*-to get insights into the influence of binding of a flat ligand on the folding landscape. To simplify the calculations we used the minimal d[GGG]_4_ construct, however, the studied strand-slippage movements are fully relevant to complete intramolecular G4s.^33, 53^ We used three structures and two positions for hemin, which together with simulations lacking hemin resulted into nine starting states, sufficiently though not completely representing the part of folding landscape represented by slip-stranded structures.

**Figure 5.**
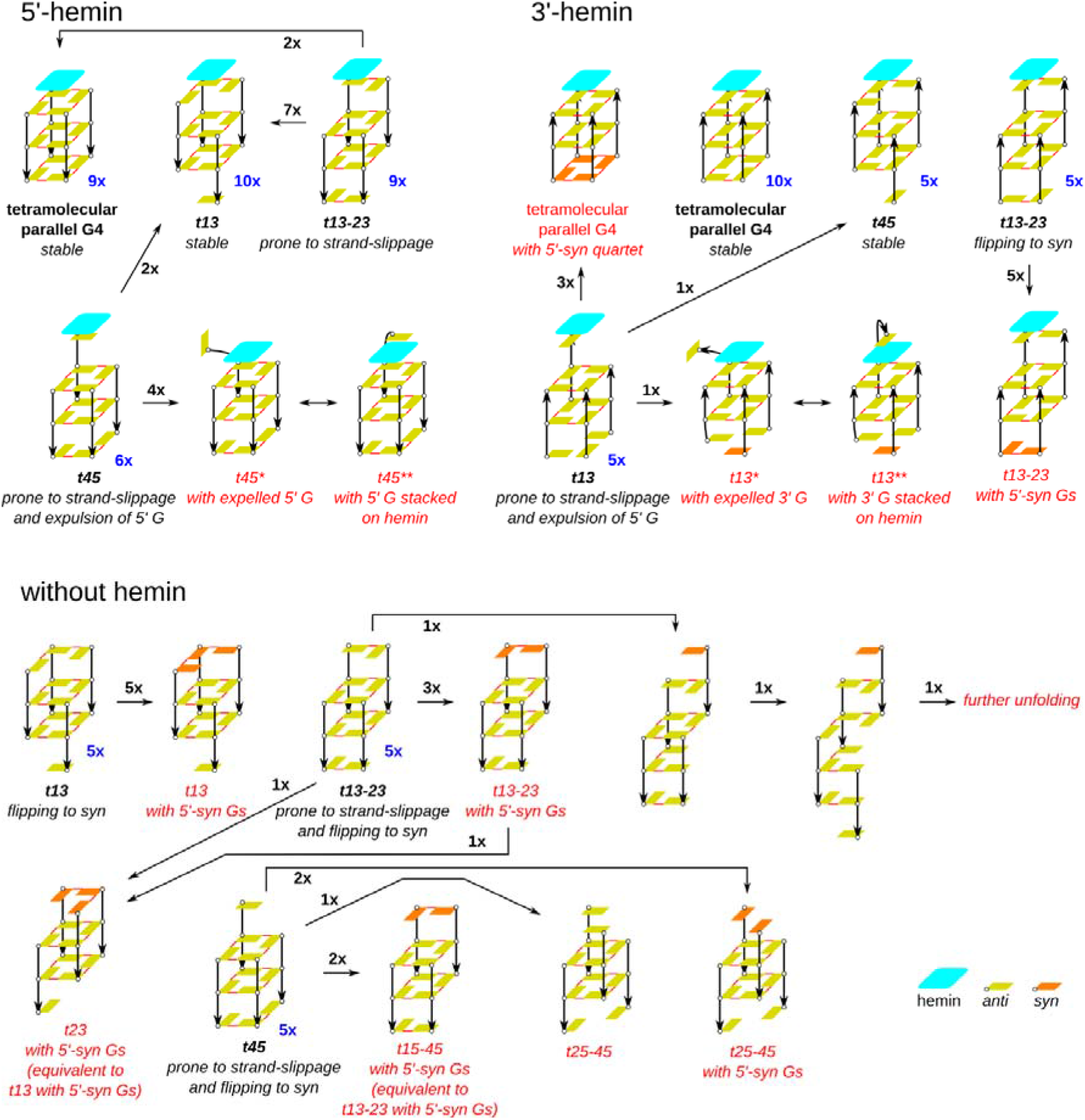
Rearrangements in complexes of tetramolecular G4s with and without hemin. Black font denotes structures used as starting structures while the red ones occurred spontaneously in simulations. Arrows show transitions observed in our simulations. The blue numbers next (bottom right) to structural schemes indicate the number of simulations initiated from a given starting structure, while the black numbers at the arrows count transitions seen in the dataset of 74 simulations. The structures marked with asterisks behave like a G4 with a flanking base near hemin. The complexes with 3□-stacked hemin are shown upside down to emphasize similarity of 5□-hemin-*t45* with 3□-hemin-*t13* and also of 5□-hemin-*t13* with 3□-hemin-*t45*. “Further unfolding” marked in this scheme is shown in Supporting Figure S12. The schematic representation is explained in the legend to Figure 2.

Development of all simulations is succinctly summarized in Figure 5. Comparison of all results reveals that the presence of hemin stabilizes slip-stranded on-pathway intermediates of folding of the parallel all-*anti* G-stem and accelerates reduction of strand-slippage towards the full three-quartet G-stem. In more details:

a. ***t13* with hemin stacked on the 5′-G-triplet:** this complex was fully stable in all five simulations with three K^+^ (at sites 5, Q, and 3). Hemin was not aligned above the *t13* central axis but shifted towards O6 atoms, and this shift was visible even in the ensemble-averaged position, as it improves stacking with the triplet (Figure 6A). The characteristic fast hemin rotation about its vertical axis is mostly stopped in *t13* by the formation of two stable H-bonds between either of the two hemin negatively-charged carboxylate groups and the Gs’ imino and amino hydrogens of the vacant Watson-Crick edge of the G-triplet (Figure 6B). The carboxylate group also provided its oxygen to the channel K^+^, replacing the oxygen atom from the missing guanine. These two H-bonds may also contribute to the shift of hemin off the G4 axis. They were occasionally disrupted, allowing hemin to rotate again until another H-bond formation occurred. The carboxylate group involved in the H-bonds served as a steric hindrance and prevented *t13* from strand slippage and formation of a complete three-quartet G4 on our simulation time scale. No exchange of the two ions present in the initial structure at sites Q and 5 with the bulk was observed, while a weak cation binding occurred at site 3. In five additional simulations starting without a cation at site 5, a cation was captured from bulk solvent in two of them; in the three other simulations, however, the site remained unoccupied. The G-triplet then adopted a cyclic triangular arrangement of the three Gs (Figure 6C), likely due to the absence of cation or a water molecule at site 5.
b. ***t13-23* with 5′-stacked hemin:** this complex was prone to strand slippage (Figure 5), which occurred in all four simulations (with three K^+^) after only a few dozens to hundreds of ns. The products were either the *t13* structure (three times) or the complete three-quartet G4 (once) formed by a simultaneous strand slippage of two strands (Supporting Movie SM2). Even two of the events leading to *t13* initiated as a double-strand slippage, but full upward movement of one of the strands was hindered by the above-described H-bonds between the hemin and a G below it. The cations at sites Q and 5 were stable in this set of simulations, while the cation at site 3 left after the simulation start. Weak cation binding at site 3 remained in only one simulation. In the set of five simulations starting with only two channel cations (at sites Q and 3), formation of *t13* was observed (four times) or the *t13-23* structure remained stable (once). The strand slippage events appeared dozens to thousands of ns after the simulation start. The Q-site cation was stable; site 5 became occupied by K^+^ after 30-300 ns and site 3 usually lost the cation upon binding of K^+^ at site 5.
c. ***t45* with 5′-stacked hemin:** this complex provided the richest dynamics among the complexes with 5′-stacked hemin (Figure 5). Hemin contacted the exposed parts of the top G-quartet within the first few ns of the five simulations with three K^+^. Then either strand slippage to *t13* occurred within hundreds of ns (twice), or the lone 5′-G was pushed out (three times) to allow hemin to fully stack on the top G-quartet, in a process that resembled the last two steps of the hemin delivery assisted by A in the 1KF1 binding simulations (Figure 3B). Before the lone G was bulged out, hemin spent a considerable time (10, 14 and 80 ns) in a tilted orientation in which it was still stacked with the top G but also in contact with two exposed Gs of the quartet below. In this position, the central iron was in vicinity of the lone G’s O6 atom (Supporting Figure S11). The dislodged lone G was then mostly moving around in the bulk solvent, but occasionally could stack on top of the hemin molecule to form a sandwich-like structure. In each of the three simulations, we observed about a dozen events in which the G was in the vicinity of the iron, which lasted up to dozens of ns. This behavior was similar to the dynamics of A in the 1KF1 TTA loops, but was more frequent. In the only simulation starting with two K^+^, we also observed expulsion of the lone G. The *t45* simulations thus revealed two contradicting trends: *i*-the route towards *t13* should accelerate the G4 folding, while *ii*-the formation of the structure in which hemin was sandwiched between the G-stem and the lone 5′-G can trap the slip-stranded structure. The exact kinetic balance between these two processes, however, cannot be elucidated from the simulations. In all simulations, only two cation binding sites were occupied, with two possible scenarios: *i*-the cation at site 5 left the G4; or *ii*-the cation at site 3 left the G4 and both remaining cations (at sites Q and 5) relocated one level downwards upon stacking of hemin to the 5′-quartet, so that the final outcome was same as in the first scenario. Infrequent (less than one per microsecond) cation exchanges with the bulk occurred at site 3.
d. ***t13* with 3′-stacked hemin:** this complex resembled that of *t45* with 5′-stacked hemin and likewise it exhibited the most diverse behavior (Figure 5). Three different scenarios were observed:

i. cations at sites 3 and 5 were expelled immediately after the start of the simulations (three times) and hemin tilted, being simultaneously stacked on the 3′-lone G and in contact with two Gs of the adjacent G-quartet. The slipped strand was (within 70-220 ns) pushed in the 5′-direction to complete the three-quartet stem. In addition, all the 5′-Gs flipped into the *syn* conformation, so that a three-quartet G4 with an all-*syn* 5′-quartet was formed, along with a cation capture from bulk to occupy site 5 (eventually becoming Q1). Such an *anti*-to-*syn* rearrangement was already observed in simulations performed with tetramolecular G4s^69^ and was explained by presence of the 5′-terminal OH groups.^81^ The *syn* preference of the 5′ - terminal Gs is due to the formation of an internal H-bond between the 5′-terminal OH and the N3 in the base.^81–83^ All *anti*-to-*syn* rearrangements reported in this study should be viewed in this context. They are typical for G4s with 5’-terminal Gs but much less likely for intramolecular G4s and for tetramolecular G4s with 5′-flanking nucleotides.
ii. a cation left site 5 immediately after the simulation start but, unlike in the previous case, hemin did not adopt the tilted position. It remained stacked on the 3′-lone G (until 240 ns) and then a double strand slippage took place (320 ns) to morph *t13* into *t45*. The movement of the fourth strand was blocked by a hemin carboxylate group that formed H-bonds with the lone 3′-G; the H-bonds remained in the newly formed G-triplet and slowed down the rotation of the hemin.
iii. the hemin tilted right after the beginning of the simulation and the 3′-lone G was expelled (210 ns). Similarly to *t45* with 5□-stacked hemin, the dislodged G either moved in the solvent or stacked on the other side of the hemin, forming a sandwich (the preferred state) in which the G could readily reach the hemin iron atom and stay there for hundreds of ns, which is considerably longer than with the *t45*. One G from the 5′-triplet flipped into the *syn* conformation.
e. ***t13-23* with 3′-stacked hemin:** this complex was structurally stable, except that the two non-quartet 5′-Gs adopted a *syn* conformation (Figure 5). The absence of slippage is consistent with the fact that vertical strand movement in G-stems with a combination of *syn* and *anti* Gs would result in a steric clash.^33^ Hemin either rotated about its vertical axis (twice) or was blocked (thrice). In the five simulations, no cation exchange with the bulk occurred at sites Q and 3, while weak cation binding with multiple exchanges was observed at site 5. The resulting dynamics thus differed from the complex of *t13-23* with hemin at the 5′-end where strand slippage occurred, because flipping of 5′-terminal Gs into *syn* was blocked by hemin.
f. ***t45* with hemin stacked on the 3′-triplet:** these simulations provided outcomes comparable to simulations performed with hemin stacked atop the 5′-triplet of *t13* (Figure 5), *i.e*., the system was stable in all five simulations. Formation of the H-bonds between the hemin carboxylate groups and the Watson-Crick edge of the G-triplet was observed, but, unlike the 5′-hemin/*t13* complex, hemin almost freely rotated around its vertical axis. Still, it was shifted away from the G4 axis (Figure 6A). Flipping of the lone 5′-G into the *syn* conformation was observed only once, because the lone G never de-stacked from the adjacent quartet in any of the simulations, therefore it did not have a chance to flip; in the only case of *syn* occurrence, it was not the base, but the sugar that turned upside down to form an internal H-bond.
g. **slip-stranded G4s without hemin**. Finally, a series of simulations was performed with a slip-stranded tetramolecular G4s without hemin (Figure 5), which are described, for space reasons, in the Supporting Information. Comparison of all results reveals that the presence of hemin stabilizes slip-stranded on-pathway intermediates of folding of the parallel all-*anti* G-stem and accelerates reduction of strand-slippage towards the full three-quartet G-stem.

**Figure 6.**
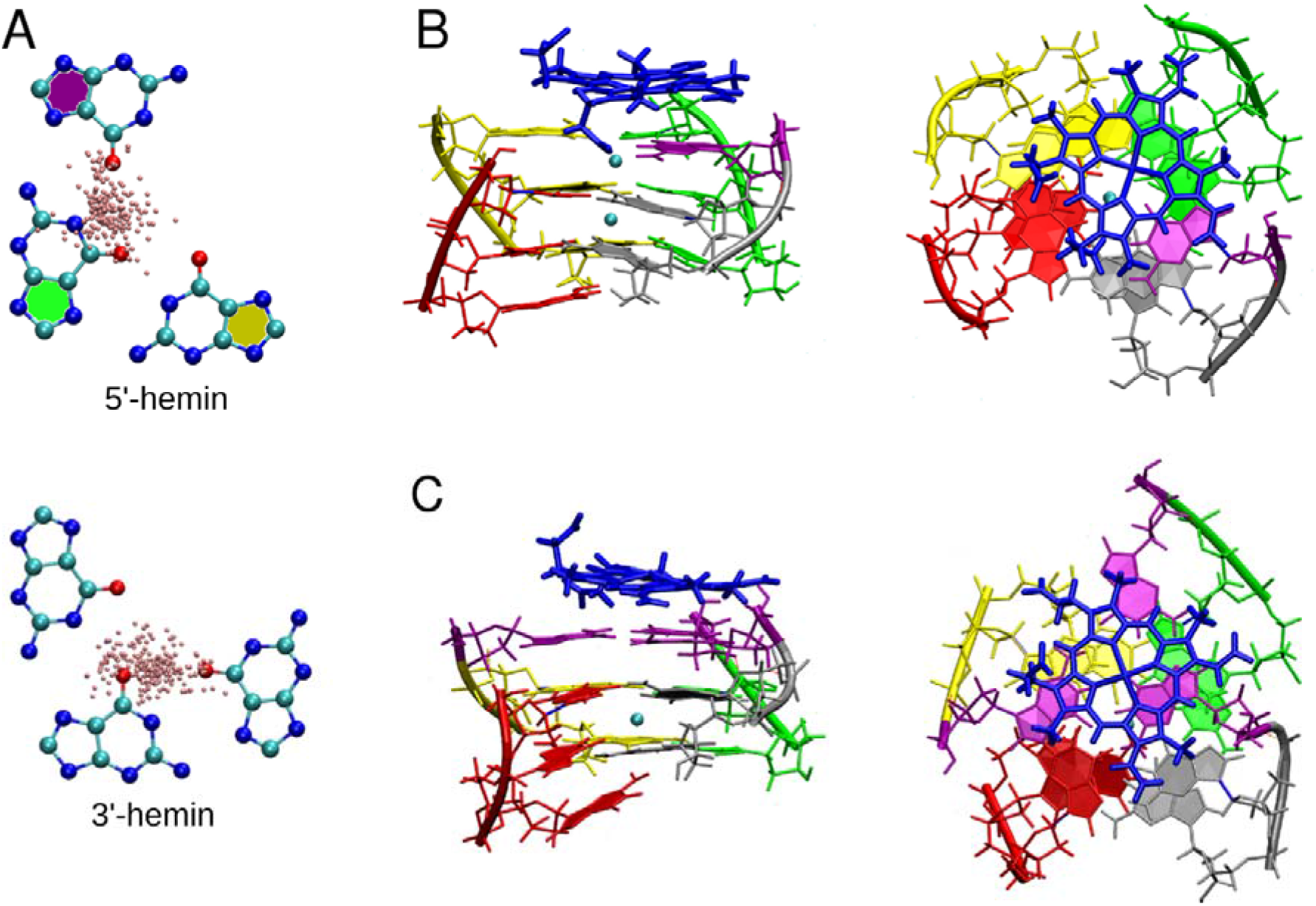
Hemin bound to the G-triplet of *t13* and *t45*. (A) 200 random positions of the iron atom (pink) above the 5□-triplet of *t13* and the 3□-triplet of *t45*. (B) 5□-hemin/*t13* complex. Two H-bonds with one of the guanines of the G-triplet beneath are formed via flexible carboxyethyl side chain. The carboxylate group also coordinates the channel cation. The ligand is visibly shifted towards the triplet, as also seen in panel A. (C) 5□-hemin/*t13* complex with the 5□-triplet under hemin in the cyclic triangular shape. In panel A, top figure, nucleotides are marked by the same color as in the structure in panel B. In panels B and C, hemin is blue while each G-strand is shown in a different color. Channel cations are cyan. In panel B, G involved in the H-bonding is shown in purple; in panel C, the three Gs forming the triangular triplet are shown in purple.

## DISCUSSION

Non-covalent association of G4s and hemin provides complexes endowed with an interesting catalytic activity that has been particularly exploited for performing peroxidase-mimicking biotransformations. Understanding the reaction mechanism is a necessary step for optimizing both the G4 scaffolds and hemin-related cofactors. Here, we have performed 483 μs of MD simulations to shed light into hemin binding to parallel-stranded G4 and its possible folding intermediates, with four interrelated goals: *i*-to decipher how hemin interacts with G4, *ii*-to gain insights into G4 dynamics, especially loop dynamics, upon hemin binding, *iii*-to assess whether hemin can affect the G4 folding landscape, mainly *via* the interaction with slip-stranded G4 folding intermediates, and *iv*-to identify which structure(s), be it a hemin/full-G4 complex or any of the G4 intermediates, could be catalytically active. It is obvious that MD simulations alone cannot describe the catalytic mechanisms. However, they can provide insights narrowing the options and suggest new possibilities.

### What is the axial ligand of hemin?

Some studies tentatively suggested that there is an axially coordinated water molecule sandwiched between heme(Fe(II)) and G4^26–29^ or cobalt(III)porphyrin and G4.^84^ However, in our opinion, the presence of the water molecule does not seem to be unambiguously proven yet for several reasons listed below. NMR signals of an alleged water molecule in heme-G4 complexes are broad, which the authors attributed to exchange of the water molecule with bulk solvent.^27^ However, while the water exchange cannot be ruled out, we think it would mean that either the complex is labile or that G4 is fragile, otherwise there is no route by which a water molecule could enter/escape from the small cavity between G4 and stacked heme. We reported here that a water molecule can be *expelled* from the channel through the quartet after manual insertion between the G4 stem and bound hemin. It occurs because of the enormous strain in the simulated structure that is unable to find any suitable position for the water molecule. The opposite process (opening the quartet and shuffling a bulk water molecule into the G-stem through a reversed pathway) is very unlikely, especially when the upper quartet is stabilized by a flat ligand. Also the signal’s integral intensity casts doubt on how many hydrogens actually correspond to the “water” signal. Unfortunately, hemin-G4 complexes evade direct NMR characterization because of the paramagnetic properties of Fe(III).^22^ Contribution from a state in which the ligand is not tightly bound to the quartet interface cannot be excluded. Raman^26^ and EPR^25^ spectroscopic analyses of hemin-G4 complexes support the transient formation of a hexa-coordinate iron, but without providing clues on the nature of the ligands. As for the NMR signals, unbound hemin could possibly contribute to the result. The structure of a heme-G4 complex with a water molecule in between hemin and G4 was also investigated theoretically by DFT calculations. However, with a distance between G4 and hemin of 3.5 Å, guanine carbonyl oxygens were suspiciously pushed away from the quartet plane by the proximal water,^27^ indicating a substantial strain (steric clash) in the computed structure. Since the QM and QM/MM calculations do not allow sampling comparable to MD simulations to sufficiently relax the starting structures, there is a risk that these calculations are performed on unrealistic (high-energy) structures.^85^ Finally, resolved crystal structures of salphen complexes stacked to 3□-quartet of a G4 do not reveal any ligand sandwiched between nickel(II) or copper(II) and G4 (PDB id’s 3QSF and 3QSC, respectively).^86^ When considering all systems discussed in this paragraph, one should also bear in mind that the results obtained with non-hemin systems (including heme(Fe(II)) instead of hemin) may not reflect properties of the hemin-G4 complex.

Collectively, the simulations performed here do not support the picture of the sandwiched water molecule, mostly because of a series of steric conflicts. No water molecule remained in the site upon hemin binding and if placed manually, it was expelled quickly. Nevertheless, we admit that fully accurate modelling of the interaction of water with iron is beyond the current limits of simulation methodology. A water molecule in proximity of the iron would become strongly polarized and its properties would change dramatically, which is not possible to capture by a non-polarizable force field (see the section “Force field limitations” in Supporting Information). Furthermore, even in case the simulations are correct and water is not a thermodynamically preferred axial ligand of hemin, catalytic activity k_cat_ of hemin-G4 complex is only about 0.1–1 s^-1^ (estimated from available kinetic data^31^), so it cannot be ruled out with a full confidence that a transiently bound water molecule between hemin and G4 could activate hemin and be responsible for its catalytic properties.

Irrespective of the simulation results, we also argue that if the iron axial ligand was a water molecule, the actual role of the G4 would be questionable, unless there is very specific, yet unknown electronic interplay in the hemin-water-G4 system. The fact that a buried water molecule would not exchange with other molecules does not make the system, in our opinion, far different from free hemin in water. Alternatively, the axial ligand could be a sterically smaller hydroxide anion, which would in many experiments be undistinguishable from ordinary water. We have performed tentative QM/MM computations and the preliminary results actually suggest that hydroxide anion is more suitable to be bound in this position than a water molecule (data not shown).

Another hypothesis found in the literature is that one of the four Gs of the G-quartet adjacent to hemin is wobbling (tilted) and provides its O6 atom as an axial ligand to the iron (Figure 1C).^30^ The simulations performed here do not support such mechanism as well. A wobbling guanine would be associated with a large energy penalty due to its partial unstacking from both the adjacent quartet and the hemin and a partial breaking of H-bonds within its G-quartet. The simulations showed that the quartet adjacent to hemin is almost perfectly flat, while without hemin, the 3′-quartet is buckled into the G4 stem (Figure S10). Recent experiments with isoguanine quartets and quintets also suggested that planar arrangement is required for reactivity.^87^ However, tilted guanines could be provided by slip-stranded G4s (see below).

We alternatively suggest that the quartet itself could be the sought-for axial ligand. Although the ensemble-averaged structure gave the expected alignment of iron atom with the channel, hemin dynamically fluctuated around the G4 axis, sampling positions closer to the G’s O6 atoms (Figure 4A). Diverse structures with shifted hemin were readily sampled in contrast to structures with wobbling G or a buried water molecule. Therefore, any of the four carbonyl oxygens could be the source of electrons for iron coordination. One could argue that there is a lack of electron density for this mechanism to be efficient, because each G’s carbonyl oxygen is involved in a hydrogen bond with a neighboring G and coordinates a K^+^ in the channel. However, the model with water sandwiched between hemin and G4 might have a similar problem. It assumes presence of two hydrogen bonds between the water and carbonyl oxygens, and thus, in fact, enough electron density from the carbonyl oxygens in the direction of hemin (Figure 1C). We also note that internal G-quartets in the G-stem contain Gs forming hydrogen bonds with neighboring Gs and coordinate *two* proximal K^+^ in the channel. This indicates that a G-quartet can coordinate chemical species both above and below its plane concomitantly.

Alternatively, the mere interaction of hemin with the electron-rich aromatic surface of the G-quartet could somehow enhance the reactivity *per se*. For example, hemin can bind to graphene and the complex was reported to enhance the peroxidase reaction about hundred times.^88–89^ Hemin bound to graphene oxide affects the peroxidation rate, too; either slightly increasing (up to about four times) or decreasing it, depending on the experimental conditions and the degree of oxidation of graphene.^90–92^ Given the similarity of graphene (oxide) surface and a G-quartet surface, both peroxidase reactions could be enhanced by the electron-rich aromatic system. Such a model is not fully in line with the existence of a hexa-coordinate iron (provided the EPR spectra reflect the actual reactive geometry) but could be rationalized by the concomitant stacking of hemin atop the quartet and coordination of iron by the four proximal carbonyl oxygens, which could act as the sixth ligand. In principle, some such stacking-caused effect could accompany the role of the other suggested ligands, such as the sandwiched water or hydroxide anion, i.e., these mechanisms are not mutually exclusive.

Our simulations further suggest that some reactive conformations could correspond to G4-folding intermediates, chiefly the slip-stranded G4s. For example, the studied *t13*, *t13-23*, and *t45* structures (Figure 2) have a G’s O6 atom in the vicinity of the iron; in the complexes with 5′-bound hemin, the angle at which the O6 interacted with the iron increased from *t13*, where hemin and G are nearly parallel (Figure 6B), to *t45*, in which hemin is tilted considerably (Figure S11). The trend was opposite in the complexes with 3′-bound hemin. Nevertheless, although such complexes may in principle be reactive (in line with the wobbling G hypothesis), they are certainly rarely populated with respect to the dominant state with G-quartet-hemin stacking. However, they could be sampled more frequently than structures of unperturbed G4 stem with wobbling G. Notably, the potential reactive conformations in slip-stranded G4s suggested by our simulations are very different from the one tentatively contemplated before (Figure 1C).^31^

Similarly, the axial ligand could be transiently provided by the loops, as the simulations show that loop bases, as well as Gs of slip-stranded intermediates, might be temporarily sandwiched between a quartet and tilted hemin (Figures 3B, 5 and S11). For the sake of completeness, stacking of hemin with a triad of slip-stranded structures may increase capability of the structure to incorporate a water molecule, though we have not seen this in the present set of simulations.

### Loops and flanking bases may bind H_2_O_2_ in the vicinity of the hemin iron atom

Catalytic activity of G4/hemin complex can be further enhanced by loop/flanking nucleotides which act as a general acid-base during the initial step of the reaction.^31, 34–36, 93^ Our simulations provide solid evidence to support this model: the three-nucleotide propeller loops of 1KF1 and single slipped Gs in *t13* and *t45* (being positioned equivalently to flanking nucleotides) are flexible enough to form a base/hemin/quartet sandwich (Figures 4B and 5). In this arrangement, the base is close to the hemin iron atom thus being suited to bind H_2_O_2_, stabilizing the transition state of the reaction and increasing its rate. The sandwich could be formed at both 5′- and 3′-ends of the G4/hemin complex. The single-nucleotide propeller loops are too short to form this type of sandwich, as demonstrated by our simulations with 2LEE. This is in line with experimental data showing no rate enhancement by short loops.^31^ This result seems to be convincingly demonstrated within the limits of the MD simulation technique.

### Hemin binding process

We initiated many of our simulations with a hemin floating in the bulk solvent and witnessed multiple binding events to G4s. In both 1KF1 and 2LEE, hemin often first bound to the G4 loops and not directly to the accessible G-quartets of the G4s. A greater versatility of intermediates was obtained with the longer loops of 1KF1 as compared to the short loops of 2LEE, but all of them were characterized by hemin stacking on bases (Figures S2 and S3). The three-nucleotide loops can adopt various conformations capable of binding hemin, stacking either to a single base or to transiently formed base pairs and even triplets. The single-nucleotide loops provide hemin with two faces of the single base only. A dynamic equilibrium makes hemin prone to detach from the loops back to the bulk solvent, or to be transferred from the loops to the 5′-quartet (Figure 3B, Supporting Movie SM1). Surprisingly, we did not observe such a hemin delivery by loop bases to the 3′-quartet, neither in 1KF1 nor 2LEE. This highlights how internal dynamics of the loops might play a major role in the binding process. It is possible (although it has not been seen in our simulations due to the affordable simulation timescale) that contacts with the loops may also facilitate the hemin unbinding events. In d[GGG]_4_, which has no loops, we have observed binding to the terminal quartets or binding to the grooves, from which hemin was usually quickly transferred to the 5′-quartet. Interestingly, rate constant of hemin binding from bulk solvent to terminal quartets was comparable for 1KF1 and d[GGG]_4_ (Tables S1 and S2), which suggests that the TTA loops do not alter the overall binding kinetics significantly. Of course, this does not necessarily apply to other loop types in other G4s, which may affect the binding differently. Spontaneous binding of a ligand to the loops, direct binding to terminal G-quartets and sliding of ligands from the loops along the backbone to the terminal G-quartets has been observed in MD simulations of G4s with other ligands, including telomestatin and BRACO19.^94–102^ A decisive loop-assisted delivery of RHPS4 to the partially exposed 3□-quartet of the human telomeric hybrid-1 type G4 has been reported, in which the propeller loop with bound ligand significantly changed its conformation and brought the ligand closer to the quartet (*Cf*. Supplementary Movie “Hybrid_G4_2HY9_02” in ref.^95^), but, to the best of our knowledge, direct loop-to-quartet ligand delivery that we observed in our study has not been reported yet. Once hemin bound to either the 5′- or 3′-quartet, either directly from the solvent or after being delivered by the loop, it stayed stacked on the G-quartet until the end of the simulations.

### 5′-*vs*. 3′-quartet hemin binding

Hemin is thought to prefer binding to the 3′-quartet of G4 with telomere-mimicking and some other sequences.^22–24, 31, 34^ It may seem contradictory to our simulations, in which ~80% preference is given to the 5′-quartet binding of the 1KF1 G4. However, our simulations are not in thermodynamic equilibrium and our time scale is insufficient to see events such as hemin unstacking from the quartets, precluding accurate estimations of the equilibrium constants. Considering the MD-estimated k_on_ in the order of 10^9^ M^-1^ s^-1^ (Table S1) and a measured K_D_ in the range of μM^31^ would lead to k_off_ in the range of 10^3^ s^-1^. It is well within the time-scales of experimental setups, thus, binding – unbinding events are definitely affecting experimental outcomes. It is therefore possible that the 5′-quartet preference is a kinetically driven process possibly enhanced by the loop conformational behavior, which brings hemin to the 5′-quartet. The 3′-quartet binding could be the thermodynamically preferred binding mode, provided that the 3′-end binding has smaller unbinding rate. Such process is orders of magnitude beyond the simulation time-scale. Furthermore, our simulations have shown that the hemin binding is a multi-pathway process and that the binding pose is an ensemble of structures differing in hemin rotation and loop nucleotides position. It would complicate a reliable quantitative calculation of binding free energies by some enhanced-sampling methods (for discussion of the limitations of enhanced-sampling methods see, *e.g*., ref.^103^).

### Hemin may increase G4-folding rates

G4-folding is a complex process, which likely involves stable off-pathway intermediates (basins), presumably misfolded G4s. These long-living kinetic traps significantly slow down the G4 folding and contribute to kinetic partitioning.^41, 44^ Under these circumstances, ligands may speed up the G4 folding process by stabilizing certain on-pathway intermediates pertinent to the native basin. In the case of the parallel-stranded G4s, substantial involvement of on-pathway structures with slipped strands has been proposed.^32–33, 53^ Flat ligands may promote their formation compared to off-pathway intermediates, reducing the kinetic partitioning, speeding up reduction of strand slippage and stabilizing the complete parallel-stranded G4. Although the full folding is far beyond the simulation time scale, our results provide indirect support to the above considerations: comparison of simulations with and without hemin showed that the ligand exerts the aforementioned effects within the folding funnel pertinent to the parallel-stranded all-*anti* G4 state (Figure 5). Strand slippage is a movement in which an entire G-stretch of a given G4 slides by one quartet level downwards or upwards;^32–33^ such movements are hindered for G4s with mixture of *syn* and *anti* Gs, thus being specific to parallel G4s. We can assume that hemin stacking stabilizes structures with the largest stacking areas, *i.e*., it favors G-quartets, and thus promotes strand slippage in the direction of complete three-quartet parallel G4s. Our simulations of slip-stranded intermediates (Figure 5) indeed showed that, in absence of hemin, strand movements commonly occur in the direction towards disruption of the structure, while in the presence of hemin (specifically when atop the 3′-quartet of *t13* and the 5′-quartet of *t13-23* and *t45*), the same slipped G4s always rearrange in the direction towards the complete G4 (Supporting Movie SM2). In addition, the time required for initiation of strand movements was shorter in simulation of slipped G4s with bound hemin. Finally, the full three-quartet d[GGG]_4_ with hemin is very rigid in simulations while without hemin it shows some volatility to strand slippage.^104^

Therefore, hemin may act as a chaperone for parallel G4s, as already demonstrated for several other G4 ligands.^105^ These results corroborate our recent investigations in which we described major stabilizing effects of hemin on parallel-stranded intramolecular G-triplexes,^45^ making them plausible folding intermediates. The simulations *per se* do not allow for drawing definite conclusions as for whether hemin shifts the equilibrium of various G4 topologies towards the parallel-stranded form, but experimental evidence suggests that porphyrinic ligands indeed favor the parallel-stranded topologies.^106–108^ Here, we propose that hemin (and similar ligands) promotes the parallel-stranded G4 folding by increasing the folding rate of the complete G4 from slip-stranded intermediates. It stabilizes these late-stage folding species and decreases their unfolding rate. These processes could stabilize folding funnel of the parallel-stranded G4 relatively to other G4 folds upon flat ligand binding. However, it should be noted that the MD simulations presented here provide neither complete nor fully accurate description of the G4 systems. Thus, limitations stemming from the nature of the force field and sampling limits must be taken into account during interpretation of the results (further discussed in Supporting Information).^41, 103, 109^ Still, the possibility that hemin binding contributes to overall stabilization of the parallel-stranded topology also by favorable interactions with its on-pathway folding intermediates seems plausible. The simulations should be accurate enough to reflect *relative* structural stabilities and rearrangement propensities of the different studied structures.

## CONCLUSIONS

We provided an atomistic description of G4/hemin interactions, describing how hemin binds to G4s and how it could affect its folding and its catalytic properties. We observed multiple routes through which hemin binds to G4s, happening either directly to solvent-exposed terminal G-quartets or *via* loop-mediated transfer to the G-quartet (Figures 3, S2 and S3, Supporting Movie SM1). Bases of three-nucleotide loops may further stack to quartet-bound hemin, forming a sandwich-like structure base/hemin/G-stem (Figure 4B).

While acknowledging the simulation limitations, we identified multiple dynamically occurring structures that could contribute to the catalytic activity of G4/hemin DNAzyme, all of which requiring directly or indirectly a G-quartet. Reaction rate enhancement could be related to hemin that is fully stacked atop terminal G-quartets of the folded parallel-stranded G4s, as thermal fluctuations of hemin often sample positions with iron atom around G’s O6 atoms, with hemin center somewhat slid off the center of the G-stem (Figure 4A). This dynamics is not apparent from ensemble-averaged data. Alternatively, mere stacking of hemin to the electron-rich aromatic surface of the G-quartet could *per se* enhance the reactivity. The model with a wobbling G is not supported by the presented simulations. They also do not support the model with a buried water molecule between hemin and G4. However, it should be cautioned that this particular outcome could be already at the edge of MD simulations reliability.

Our results also offer new insights how loops and flanking bases could enhance the rate of peroxidase reaction *via* the transient formation of base/hemin/G4 sandwich structures (Figure 4B), which supports the model in which the base can interact with H_2_O_2_ in the vicinity of the hemin iron atom and promote the proton transfer. We finally show that hemin may facilitate the folding of parallel-stranded G4s from slip-stranded G4 intermediates, driving strand slippage in the direction of a complete G4 (Figure 5; Supporting Movie SM2), thus acting as a G4 chaperone. This might have important consequences *in vivo* as G4/hemin interaction is likely to occur in cells. We believe that the characteristics of the G4/hemin systems described here are intrinsic properties that could guide further experimental studies.

## ACKNOWLEDGMENTS

This work was supported by the project SYMBIT [CZ.02.1.01/0.0/0.0/15_003/0000477] financed by the European Regional Development Fund with financial contribution from the MEYS CR; the work was also supported by the Czech Science Foundation [21-23718S]. J.S. acknowledges support by Praemium Academiae; P.S. greatly appreciates access to CESNET storage facilities provided by the project “e-INFRA CZ” under the program “Projects of Large Research, Development, and Innovations Infrastructures” [LM2018140]; M.O. acknowledges the Faculty of Science, Palacký University, Olomouc for support; J.Z. acknowledges the financial support of the National Natural Science Foundation of China [21977045]; and Fundamental Research Funds for the Central Universities [02051430210]; J.L.M. acknowledges the funds of Nanjing University [020514912216]; and D.M. acknowledges the Agence Nationale de la Recherche [ANR-18-CE07-0017-03]; and the European Union [PO FEDER-FSE Bourgogne 2014/2020 programs].

## SUPPORTING INFORMATION

Supporting Information consists of the following files: Supporting_information.pdf contains description of equilibration protocol, detailed results of MD simulations of slipped tetramolecular G4s without hemin, additional discussion of force field limitations, all Supporting Figures (S1–S14) and Tables (S1 and S2); Supporting_movie_SM1.avi is a 150 ns long portion of trajectory showing transfer of hemin to the 5□-terminal quartet of 1KF1 facilitated by a loop adenine; Supporting_movie_SM2.avi is a 60 ns long portion of trajectory showing strand slippage of a slipped tetramolecular G4 into a full three-quartet G4 facilitated by stacked hemin.

