## Supporting Information for "Insights into G-Quadruplex–Hemin Dynamics Using Atomistic Simulations: Implications for Reactivity and Folding"

[1] Institute of Biophysics of the Czech Academy of Sciences, Královopolská 135, 612 65 Brno, Czech Republic.
[2] Regional Centre of Advanced Technologies and Materials, Czech Advanced Technology and Research Institute, Palacký University, Křížkovského 8, 779 00 Olomouc, Czech Republic.
[3] State Key Laboratory of Analytical Chemistry for Life Science, School of Chemistry & Chemical Engineering, Nanjing University, Nanjing 210023, China.
[4] Institut de Chimie Moléculaire (ICMUB), CNRS UMR6302, UBFC Dijon, 21078, France.
[5] Laboratoire d'Optique et Biosciences, Ecole Polytechnique, CNRS, INSERM, Institut Polytechnique de Paris, 91128 Palaiseau cedex, France.
[§] Present address: Department of Pharmacy, School of Applied Sciences, University of Huddersfield, Huddersfield HD1 3DH, United Kingdom
* corresponding authors

### SUPPORTING METHODS

**Equilibration protocol**. The equilibration before MD simulations starts with 500 steps of steepest descent minimisation followed by 500 steps of conjugate gradient minimisation with 25 kcal mol$^{-1}$ A$^{-2}$ position restraints on DNA atoms. Then the system is heated from 0 to 300 K during 100 ps with position restraints of 25 kcal mol$^{-1}$ A$^{-2}$ on DNA and with constant volume. Afterwards, the system undergoes minimisation with 5 kcal mol$^{-1}$ A$^{-2}$ restraints on DNA atoms using 500 steps of steepest descent method followed by 500 steps using conjugate gradient. Continuing in position restraints on DNA atoms of 5 kcal mol$^{-1}$ A$^{-2}$, the system is equilibrated during 50 ps at a constant temperature of 300 K and pressure of 1 bar. Then, an analogous series of alternating minimisations and equilibrations is performed, consecutively using decreasing position restraints of 4, 3, 2 and 1 kcal mol$^{-1}$ A$^{-2}$. Finally, an equilibration using position restraints of 0.5 kcal mol$^{-1}$ A$^{-2}$ and starting velocities from the previous equilibration step, followed by a short 50 ps free molecular dynamics takes place. Temperature and pressure coupling constant used during equilibration is set to 0.2 ps, while coupling during the last molecular dynamics phase is set to 5 ps.

### SUPPORTING RESULTS

**Slipped G-quadruplex without hemin**. For a comparison with simulations of slip-stranded structures with bound hemin, we have carried out a series of simulations of slipped tetramolecular G4 structures without hemin. Most importantly, they showed that the tendency to transform in the direction of full G4 was higher in complexes with hemin. The simulations thus support the view that the presence of hemin stabilizes strand-slipped intermediates of a parallel G-stem and may accelerate the reduction of strand-slippage towards the full three-quartet stem.

The *t13* structure without hemin was stable in all five simulations and no significant strand dynamics was observed (Main text Figure 5). However, guanines in the 5′-triplet had a clear tendency to flip from the *anti* conformation to *syn*; in two simulations all three guanines flipped and formed a *syn* G-triplet, while in three the transition was partial with one guanine retaining the *anti* conformation; see the discussion in the main text explaining the tendency to *anti* to *syn* flips for 5′-

terminal Gs. The ion at site Q remained bound whereas the ion at site 5 was exchanged with the bulk on average once per five microseconds.

The *t13-23* structure without hemin remained intact with no cation exchange at site Q in two simulations (Figure 5). In two other simulations, it underwent a strand slippage to form the *t23* structure (equivalent to *t13*). The cation bound at site Q of the original *t13-23* two-quartet G4 remained at its place even during the transition and stayed bound in the channel of the newly formed *t23* structure. As in the simulations of *t13*, guanines at 5′-end that were not part of a quartet tended to flip to the *syn* conformation. In these four simulations we observed a visible trend to bind a cation at site 5, but not at site 3. In another simulation, the structure fully unfolded (Figure 5, Supporting Figure S12). The process began at 0.65 μs by a strand slippage in the 3′-direction of a strand that already was slipped in this direction in the starting structure. A structure with only one quartet was thus formed. At 0.70 μs, the neighboring strand that was not slipped in the starting structure slipped in the 5′-direction, so the only remaining quartet was lost and the structure quickly collapsed into an assembly of four strands without any G:G base pairs typical for G4s.

The *t45* structure without hemin always transformed into other structures (Figure 5). In all five simulations, one strand slipped one step upwards, so a two-quartet G4 still remained. From these, in two simulations the slipped strands were neighboring, and thus *t15-45* (equivalent to *t13-23*) was formed. Surprisingly, in one of them, 10 ns before the mentioned strand slippage event, another strand slipped, but downwards, so only one quartet was left. This was immediately followed by a counter-strand slippage upwards, so two quartets restored fast, preventing the collapse of G4. In the other three simulations the opposing (non-neighboring) strands slipped, resulting in the *t25-45* structure. Two guanines on either side of the G4 core, be it above or below, could either form a G:G base pair (Supporting Figure S13), or their O6 and N7 atoms interacted with a cation if it was there. Regardless of the outcome, guanines at the 5′-end had a tendency to flip to the *syn* conformation. The cation at site Q did not move upon the strand slippage and remained bound inside the two-quartet G4 core in all the simulations. Cations bound at sites 5 and 3 were exchanged with the bulk several times over the course of the simulations. The arrival of a cation at site 5 or 3 resulted in expulsion of a cation from the other site so that at most two ions were bound at the same time.

### SUPPORTING DISCUSSION

**Force field limitations.** Past MD simulation studies provided important insights into G4 structural dynamics, demonstrating that MD simulations represent a valid complement to diverse experimental investigations. However, when assessing the primary MD simulation data, it is always important to take into consideration sampling limitations, and also force-field accuracy limitations. In general, the major limitations of the current nucleic acid force fields affecting G4 simulations are under-stabilization and isotropic nature of quartet's H-bonds,[1-4] under-stabilization of cation-quartet interactions,[5-6] potential over-stabilization of base stacking[7-9] and evidently excessive mutual repulsion of cations inside the channel.[5, 10-12] However, these limitations should not affect the interpretation of our results, as we have taken them into consideration when assessing the simulation outcomes.

Isotropic nature (i.e., lack of an explicity directionality term) of hydrogen bonds rather commonly causes formation of quartets with bifurcated hydrogen bonding, i.e., N7 binds to both N2 amino group and N1 imino group.[11] Weaker hydrogen bonds make various G4-related structures, e.g., G-triplexes and G-hairpins, less stable than they in fact are.[1-2] A remedy for both the deficiencies in a form of an additional potential supporting hydrogen bonds, called HBfix[3] and its generalized form, gHBfix,[2] has been suggested. Alternative approach targeting the van der Waals combination rules has been suggested, too.[4] Nevertheless, none of the approaches is perfect and future improvements of the force field, for example, the introduction of bond directionality, could help further. Fortunately, it is expected that underestimated stability of the hydrogen bonds is somewhat compensated for by sticky bases in G4 stems (see the next paragraph), so the (g)HBfix potential was not required in our calculations. The simulations were sufficiently stable on the affordable simulation time scale.

Over-stabilization of base-stacking simply makes bases stickier. Strong stacking is probably partly responsible for non-ideal conformational behavior of propeller loops, though the origin of problems in the description of propeller loops are not fully clear.[7-9] The TTA loops of 1KF1 tend to form rigid triple-stack structures in MD simulations while they also do not reproduce the γ-trans backbone states seen in many X-ray structures.[8] Formation of the spurious over-stacked loop conformation, however, seems to be reduced by hemin stacked to the loops, so while we cannot guarantee perfect conformational behavior and correct populations of loop substates, the loops with bound hemin appear to be flexible enough and were not stuck in a single conformation. The misbehavior of the propeller loops may be more significant when hemin is already bound to the terminal quartets. It may lead to underestimation of the rate at which the loop bases move to form the sandwich structures with hemin though, conversely, life-time of these structures could be overestimated. Thus, imperfect behavior of propeller loops should not affect the qualitative interpretation of our data. Similarly, over-stabilized stacking could increase life-times (decreased dissociation rates) of hemin-G4 complexes. This has literally no practical impact on stacking of hemin to terminal quartets, for which even ten times decreased life-times, should stacking be weaker, would still result in life-times much longer than the simulation time scale. Stronger stacking could have minor impact on the dissociation rate of hemin from loops in the simulations. Nevertheless, should stacking be weaker, increased dissociation rate of loop-hemin complexes would only be partly beneficial for increased association rate of hemin with terminal quartets, because freed loops would again act as competitive binding sites for hemin. Altogether, we suppose that the over-stabilization of stacking has lesser impact on our estimation of kinetic constant values than the uncertainty coming from the limited simulation sampling.

Description of cation binding to bases, including those bound to G-quartets, suffers from the lack of polarization in the force field. As a result, binding of such cations to bases is weaker than it should be.[5-6] The overall effect is a combination of expected errors stemming from the lack of reorganization of electron density in Gs upon the formation of a G-quartet (with the bound ion) and no electronic-structure response of water molecules that are around cations.

In addition, a G4-characteristic force-field flaw is excessive inter-cation repulsion and it is relevant when two or more cations are bound in the G4 channel. It has been shown that positive charges of neighboring cations are not (electronically) shielded enough by the carbonyl oxygens due to, again, the lack of polarization in the force field.[5, 10-11] In other words, the force field describes the cations in the channel trivially as Lennard-Jones spheres with fixed +1 e point charge and the inter-cation repulsion is then exaggerated. In reality, the 'apparent' or effective cation charges that would appropriately describe the ion-ion interaction are certainly reduced by polarization of the G-quartets by the ions, in a highly geometry-dependent manner, *i.e.*, the balance cannot be captured by fixed-charge models. There is an intricate polarization communication between all the ions and all the G-quartets. These many-body effects are by definition zeroed in all pair-additive force fields. The resultant excessive inter-cation repulsion is cumulative, appears first with two ions in the channel and is pronounced in G4s with three or more cations.[12] In the present work, the systems possibly affected by this artifact are the slipped G4s, *i.e.*, *t13*, *t13-23* and *t45* structures. When starting with three cations in the channel of these structures, the bottom one was immediately expelled. The speed of this process can likely be attributed to excessive inter-cation repulsion. On the other hand, the repulsion might not necessarily be the only factor in play; the bottom channel site is not complete, because it is between a quartet and a lone guanine/G:G base pair/G-triplet. Therefore the binding site might be weak or even non-existent even in reality. Anyway, given that hemin was placed above the studied structures it does not seem that the cation leaving the bottom site of the slipped structures affected the simulation outcome in any significant way. Most importantly, these problems should not affect the **relative difference** in behavior of structures with and without bound hemin. Regarding ion distribution, polarizable force fields should have an advantage.[5-6, 13] Of note, in the simulations, iron(III) of hemin is not present as a cation with a charge of +3 e, but instead partial charges of all hemin atoms are parametrized as a whole group. It means that electron distribution (electrostatic potential to be more precise) observed in QM is transferred to MD to some degree. Thus, the iron in our simulations has a charge of +0.2492 e, which eliminates potential excessive repulsion between iron(III) and the nearby channel $K^+$ at sites Q1 and Q2. We have verified this in a large-box simulation of 1KF1 by comparison of the average position of sites Q1 and Q2 before and after hemin binding. In both cases, position of sites Q1 and Q2 relative to the nearby guanines was virtually identical, which confirms no excessive inter-cation repulsion caused by hemin (Figure S14).

A specific uncertainty in our simulations is connected to the lack of polarization and electronic structure effects, which limits the description of the iron(III) species. In this work we have used hemin parameters developed by Shahrokh et al.[14] for ferric penta-coordinate high-spin hemin bound in a cytochrome P450, *i.e.*, not in bulk solvent and its native axial ligand is cysteine. This choice may raise questions about the validity of the description of hemin. To shed more light on this topic, we have later developed our own set of parameters for high-spin hexa-coordinate hemin and studied its association and interaction with a four-quartet G4 (preliminary data, not shown). It has turned out that the results regarding hemin-G4 interaction presented in this paper are essentially identical to those we have obtained with our new parameters. In addition, we have found that when we reduced the iron(III) radius from 1.8 to 1.3 Å (with appropriate adjustment of RESP charges; this radius has been considered for hexavalent heme derivatives by Shahrokh et al.[14]), residency time of a water molecule placed between hemin and G4 has increased at least one fold, but it still does not get trapped there upon the hemin binding process, nor does it enter the site later on during the course of simulations. In other words, the presented results are robust with respect to hemin parameter changes within the limits of the pair-additive force field approximation.

However, in reality, electronic structure of a water molecule between hemin and G4 and any water molecule in the vicinity of the iron(III) on the other side of hemin would be affected by the interaction with the iron, and therefore quite different from ordinary water molecules.[15] It can be affected by electronic-structure effects which are not reliably includable in the simple force-field description. The water properties could be affected by the suspected binding to the quartet, too. Despite that there is not much steric volume to accommodate a water molecule between stacked hemin and G-quartet, as seen in the MD simulations, such electronic structure effects can hypothetically enhance its binding capability. We have presently no data to assess how such effects (absent in the simulations) may affect the results.

In addition, our recent preliminary simulations of water molecules inserted into G4 stems suggest that behavior of water molecules in the channel is moderately water-model-dependent. Water molecules of some water models show some capability to stay in the channel while others tend to be expelled from the channel, including SPC/E used in this study.[12] We have also evidenced transient insertions of water molecules (from inside the channel) into G-quartets. However, it is presently impossible to decide which water model is more realistic, due to lack of any experimental information that could serve as a benchmark. Still, the differences among the water models are not large enough to qualitatively change the basic results (inability to stably host a water between hemin and G4), to our opinion.

Nevertheless, considering all uncertainties, although the present simulations suggest that stabilization of a water molecule between the hemin and the stem is unlikely due to a major steric conflict, *description of this specific interaction is potentially affected by the largest force-field uncertainty* among all the interactions analyzed in the paper.

The force-field limitations are also one of the reasons why we did not attempt any enhanced-sampling simulations to make quantitative free-energy predictions. The other reason is that with our simulation time-scale, we have been capable to see sufficient number of structural transitions in unbiased simulations to draw the basic conclusions and predictions of our study. In other words, we suggest that the use of enhanced-sampling methods was not necessary in this specific work. Enhanced-sampling methods are not a panacea and, for example, application of approaches based on collective variables (CV) could hide some movements that are not specifically accelerated by the chosen CV, and thus distort the picture.

**SUPPORTING TABLES.**

**Table S1**. Estimation of the rate of the hemin-1KF1 complex formation from large-box simulations.[a]

| 1KF1 simulation | Hemin and G4 concentration *c* (M) | Time for binding to quartets *t* (s) | Binding rate $k_{on}$ ($M^{-1}$ $s^{-1}$) |
|---|---|---|---|
| Sim. 1a | $4.61 \times 10^{-4}$ | $2.26 \times 10^{-6}$ | $0.96 \times 10^{9}$ |
| Sim. 2a | $4.08 \times 10^{-4}$ | $1.20 \times 10^{-6}$ | $2.04 \times 10^{9}$ |
| Sim. 3a | $4.40 \times 10^{-4}$ | $1.24 \times 10^{-6}$ | $1.83 \times 10^{9}$ |
| Sim. 1b[b] | $4.61 \times 10^{-4}$ | $3.70 \times 10^{-6}$ | $0.66 \times 10^{9}$ |
| Sim. 2b[b] | $4.08 \times 10^{-4}$ | $1.12 \times 10^{-6}$ | $2.03 \times 10^{9}$ |
| Sim. 3b[b] | $4.40 \times 10^{-4}$ | $0.33 \times 10^{-6}$ | $6.61 \times 10^{9}$ |

[a] Although six simulations do not provide statistically converged results, they are sufficient for a rough estimate (binding time observed in individual simulations was used as an estimator of second-order reaction half-life), calculated by the formula $k_{on} = 1/c/t$, that the $k_{on}$ is in the range of $10^9$ $M^{-1}$ $s^{-1.}$ This is consistent with values obtained from the midsize-box simulations (Table S2).

[b] Exactly the same setup as in the 1a-2a-3a simulation set, but using Langevin thermostat with Monte Carlo barostat

**Table S2**. Estimation of the rate of the hemin-tetramolecular d[GGG]$_4$ G4 complex formation from midsize-box simulations.[a]

| d[GGG]$_4$ simulation[b] | Hemin and G4 concentration *c* (M) | Time for binding to quartets *t* (s) | Binding rate $k_{on}$ ($M^{-1}$ $s^{-1}$) |
|---|---|---|---|
| Sim. 4 | $9.77 \times 10^{-3}$ | $13.2 \times 10^{-8}$ | $0.78 \times 10^{9}$ |
| Sim. 5 | $5.81 \times 10^{-3}$ | $1.9 \times 10^{-8}$ | $9.06 \times 10^{9}$ |
| Sim. 6 | $5.87 \times 10^{-3}$ | $6.6 \times 10^{-8}$ | $2.58 \times 10^{9}$ |
| Sim. 7 | $7.72 \times 10^{-3}$ | $6.7 \times 10^{-8}$ | $1.93 \times 10^{9}$ |
| Sim. 8 | $5.48 \times 10^{-3}$ | $3.0 \times 10^{-8}$ | $6.08 \times 10^{9}$ |
| Sim. 9 | $5.89 \times 10^{-3}$ | $7.9 \times 10^{-8}$ | $2.15 \times 10^{9}$ |
| Sim. 10 | $7.22 \times 10^{-3}$ | $3.6 \times 10^{-8}$ | $3.85 \times 10^{9}$ |
| Sim. 11 | $6.89 \times 10^{-3}$ | $2.1 \times 10^{-8}$ | $6.91 \times 10^{9}$ |
| Sim. 12 | $9.07 \times 10^{-3}$ | $0.3 \times 10^{-8}$ | $36.73 \times 10^{9}$ |
| Sim. 13 | $7.41 \times 10^{-3}$ | $3.2 \times 10^{-8}$ | $4.21 \times 10^{9}$ |

[a] As for the large-box simulation (Table S1) a rough estimate of $k_{on}$ (binding time observed in individual simulations was used as an estimator of second-order reaction half-life), calculated by the formula $k_{on} = 1/c/t$, gives the value in the range of $10^9$ $M^{-1}$ $s^{-1}$

[b] All simulations in this set used Langevin thermostat with Monte Carlo barostat

### SUPPORTING FIGURES

Figures are listed in the order they appear in the main text. Figures S4 and S5 are divided into three and two parts, respectively. The legend to them is below their first part.

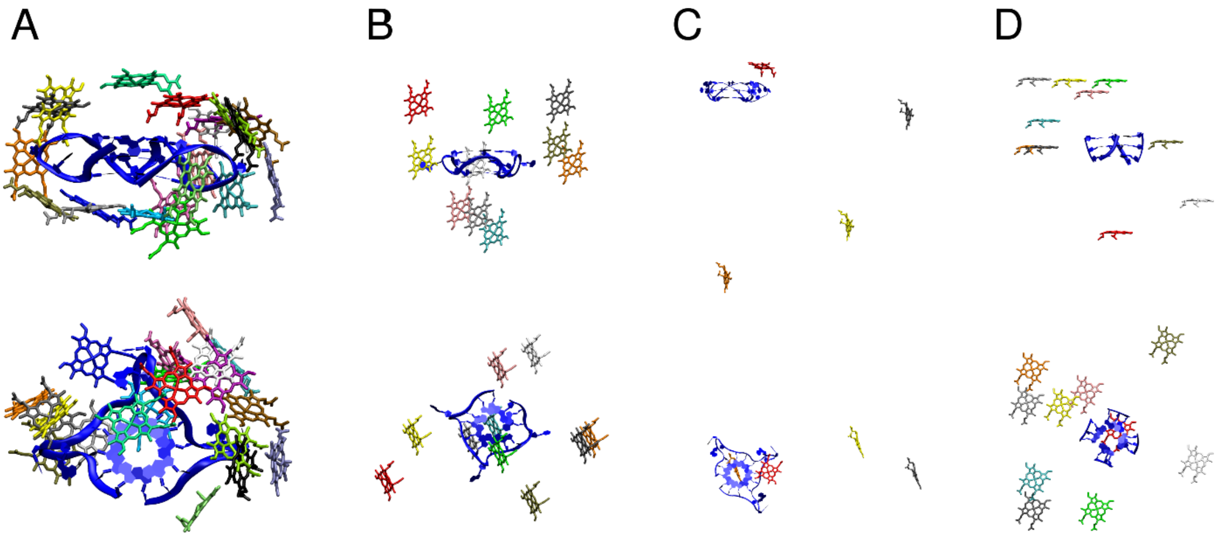

**Figure S1**. Starting position of hemin with respect to G4 in the simulations. (A) Twenty hemin positions around 1KF1; (B) ten hemin positions around 2LEE; (C) three positions of hemin used for six large-box simulations, each position was simulated twice (once using weak-coupling thermostat and barostat and once using Langevin thermostat and Monte-Carlo barostat). In these six simulations, hemin was starting far away from the G4, with a distance of about 80 Å from G4 (measured using their centers of mass). The red hemin in vicinity of G4 comes from the panel A and is used for visual comparison; (D) ten positions of hemin around tetramolecular three-quartet d[GGG]$_4$ G4 used for midsize-box simulations. G4 (either 1KF1 or 2LEE or d[GGG]$_4$) is shown in blue, each different hemin starting position is depicted by a different color. In all 46 binding simulations, the hemin finally stacked on one of the terminal G-quartets.

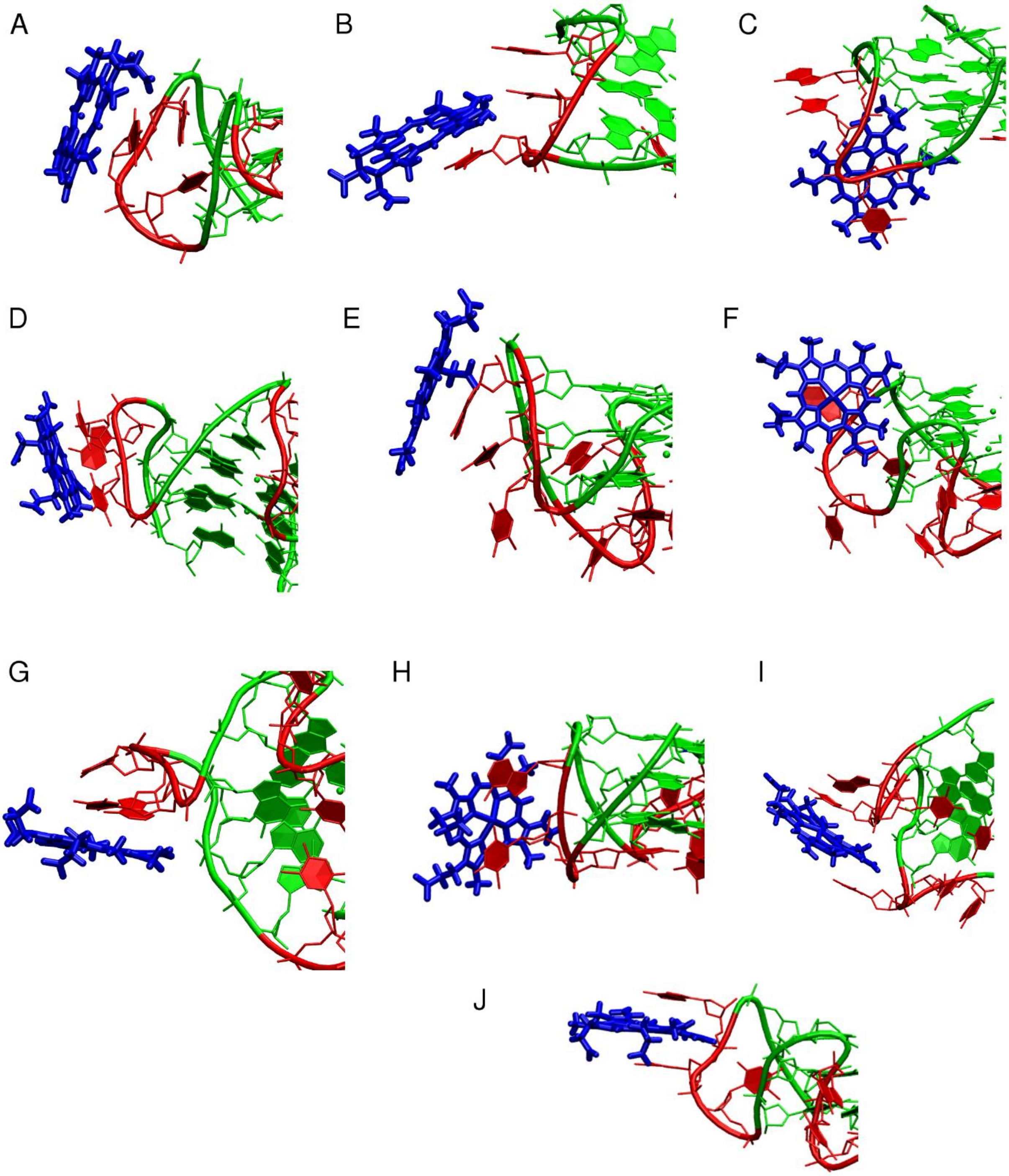

**Figure S2**. Hemin binding to the loops of 1KF1. (A) stacking to a single T, (B) stacking to a single T and side chain interaction with another T, (C) stacking to a single T and insertion to the groove, (D) stacking to a T:T base pair and side chain interaction with A, (E) stacking to A, (F) stacking to A and insertion to the groove, (G) stacking to a T:A base pair, (H) another stacking to a T:A base pair, (I) stacking sandwich T:hemin:T between two loops, (J) hemin intercalated in between two consecutive loop bases. Hemin is shown in blue, loops are red and G-stem is green; only the relevant part of the hemin-G4 complex is shown.

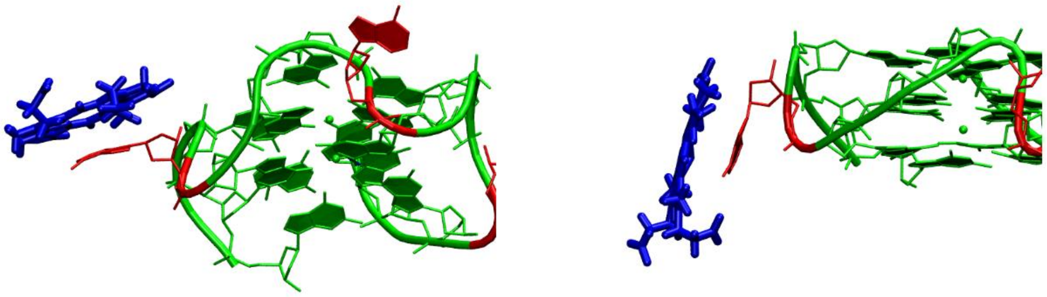

**Figure S3**. Hemin binding to the loops of 2LEE. Two positions of hemin stacked to A (no binding to C was seen). Hemin is shown in blue, loops are red and G-stem is green; only the relevant part of the hemin-G4 complex is shown.

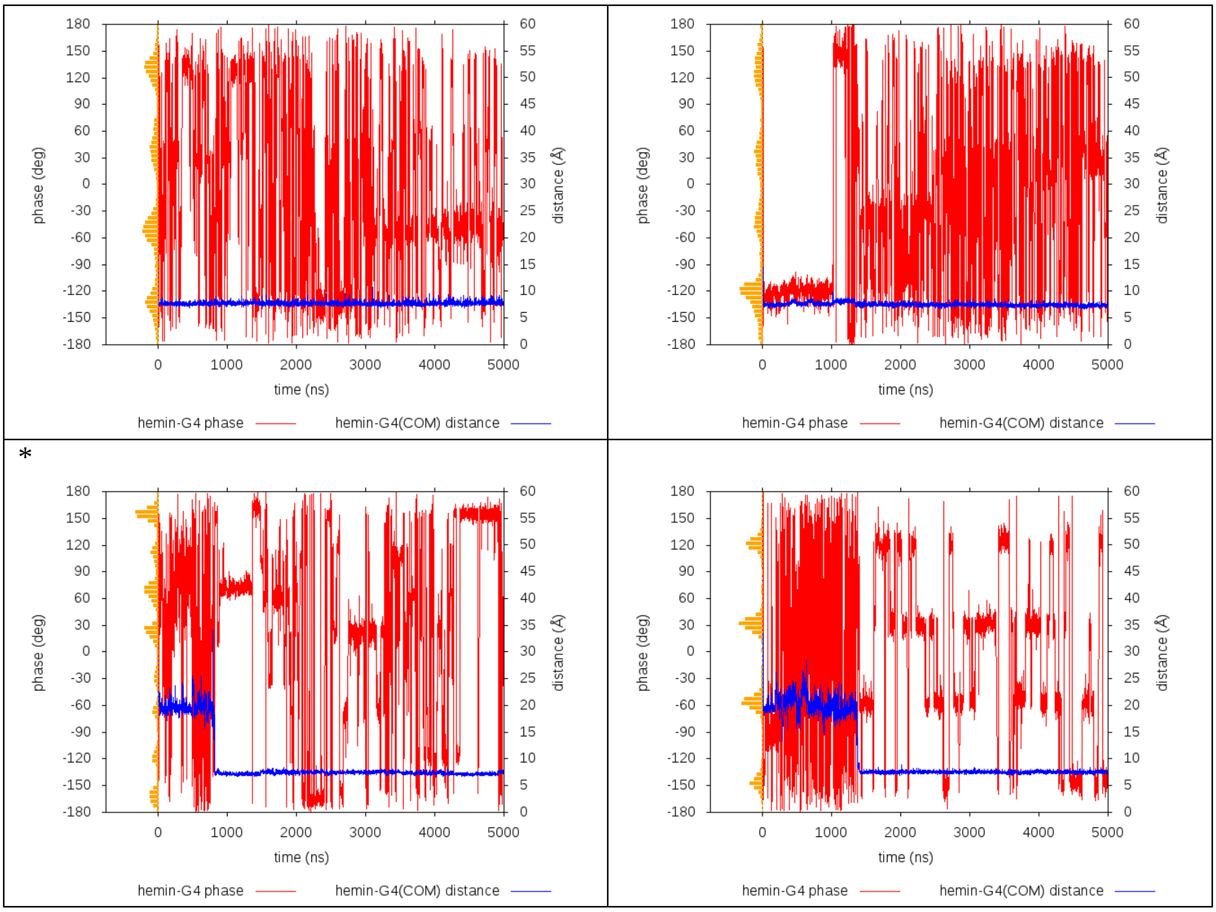

**Figure S4 (part 1/3)**. Hemin binding to 1KF1 G4 and its rotation in twenty simulations. The distance between hemin and G4 center of mass (COM) (blue line) indicates whether hemin already binds. Plateaus indicate bound states (on loops or quartets), while fluctuations mean freely moving hemin. It can be seen that once hemin bound to the quartet (the shortest possible distance below 8 Å), it never unbound. Simulations with hemin stacked to the 3′-quartet are marked by ‘*’ at the upper left corner, otherwise hemin was stacked to the 5′-quartet. The phase is an arbitrarily chosen dihedral angle defined by the following four points: 1) COM of C1′ atoms of G2 and G20 (i.e. Gs of the middle quartet from the first and last G-stretch), 2) COM of base heavy atoms of guanines forming the middle quartet, 3) hemin Fe(III), 4) hemin ring hydrogen atom located between the two carboxyethyl side chains. It characterizes the rotation of hemin about the vertical axis. The number is meaningful only after hemin is bound. As can be seen from the graphs, the rate of rotation varies and is affected by transient presence of interactions of hemin side chains with G4 backbone and loops. The orange histogram in the left part indicates the distribution of the phase dihedral angle for the relevant portion of the trajectory. See the main text Figure 3C for phase distribution summed over all simulations, Figures S5 and S6 for details of 2LEE G4, and Figures S7, S8 and S9 for details of the tetramolecular G4.

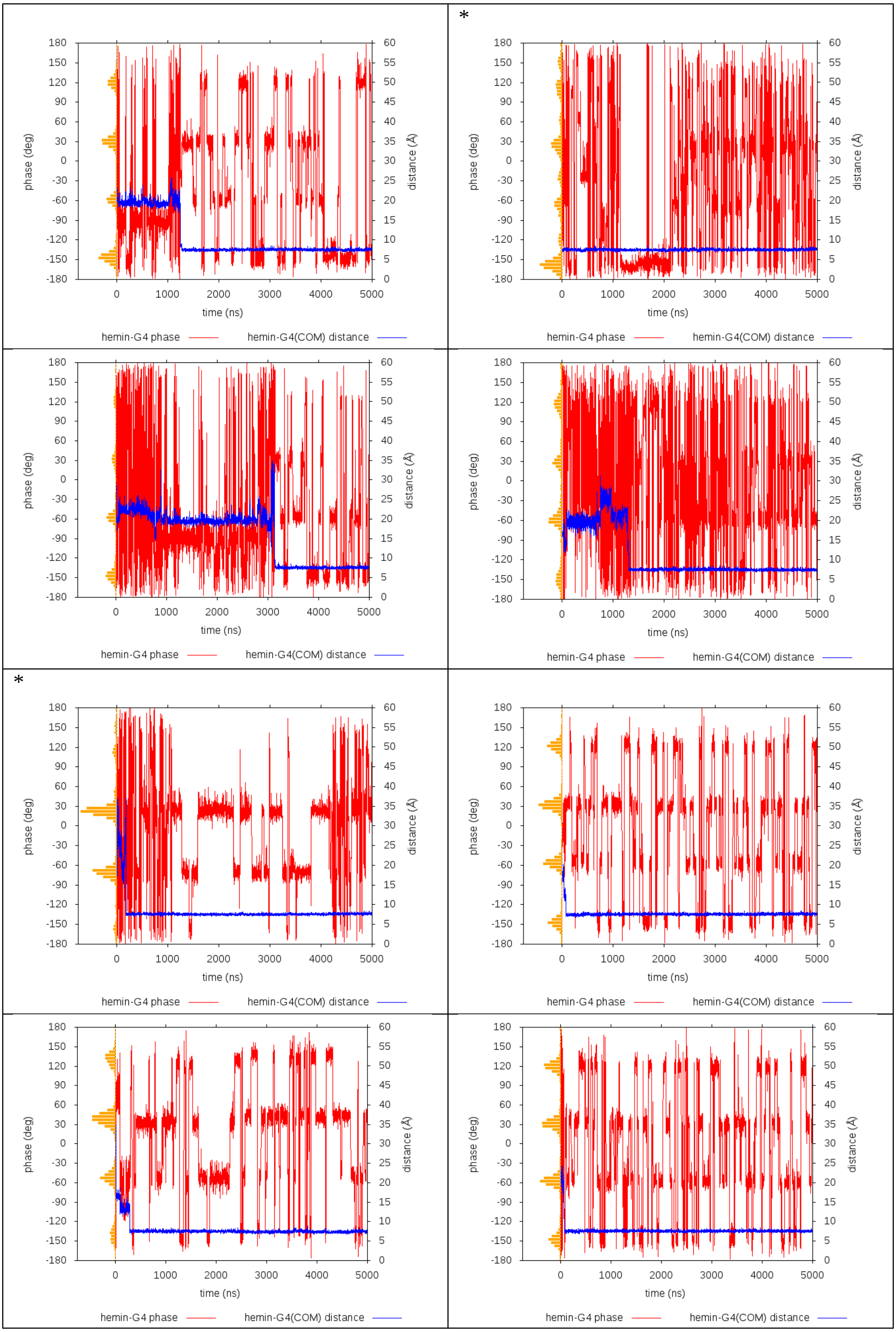

**Figure S4 (part 2/3)**. Hemin binding to 1KF1 G4 and its rotation. See the first part for full explanation.

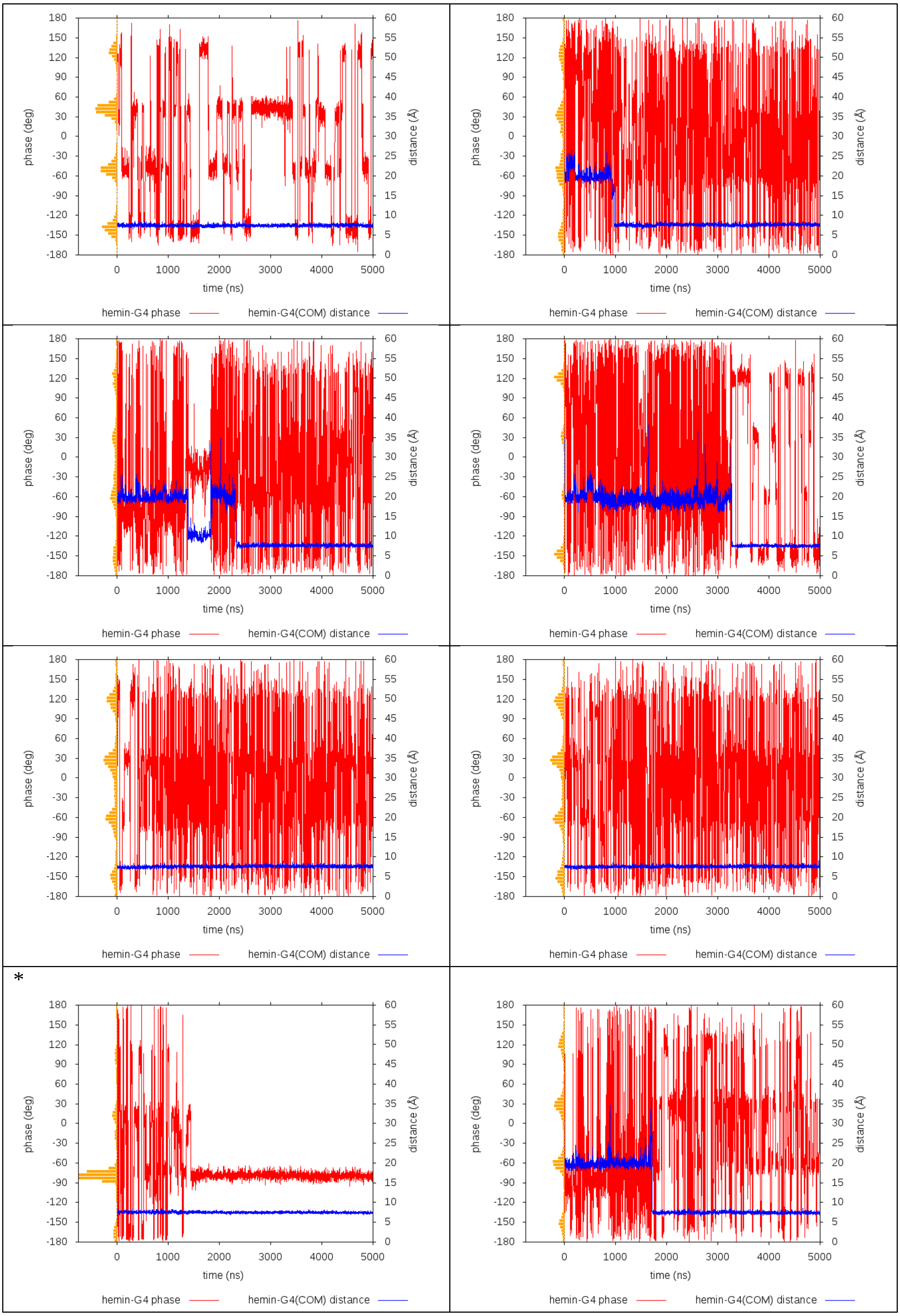

**Figure S4 (part 3/3)**. Hemin binding to 1KF1 G4 and its rotation. See the first part for full explanation.

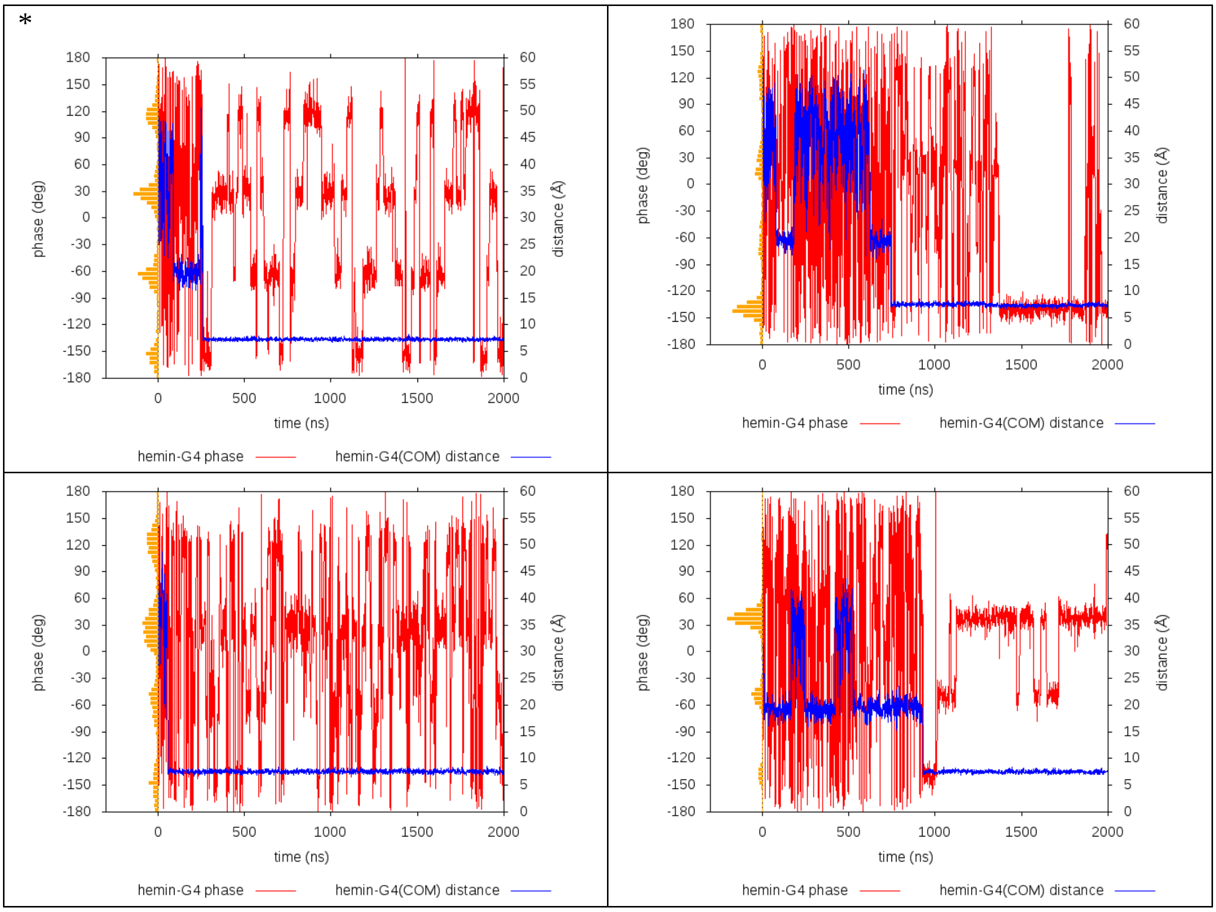

**Figure S5 (part 1/2)**. Hemin binding to 2LEE G4 and its rotation in ten simulations. Like for 1KF1 (Figure S4), the distance between hemin and G4 COM (blue line) indicates whether hemin already binds. Plateaus indicate bound states (on loops or quartets), while fluctuations mean freely moving hemin. It can be seen that once hemin bound to the quartet (the shortest possible distance below 8 Å), it never unbound. Simulations with hemin stacked to the 3′-quartet are marked by '*' at the upper left corner, otherwise hemin was stacked to the 5′-quartet. The phase is an arbitrarily chosen dihedral angle defined by the following four points: 1) COM of C1′ atoms of G2 and G14 (i.e. Gs of the middle quartet from the first and last G-stretch), 2) COM of base heavy atoms of guanines forming the middle quartet, 3) hemin Fe(III), 4) hemin ring hydrogen atom located between the two carboxyethyl side chains. It characterizes the rotation of hemin about the vertical axis. The number is meaningful only since hemin is bound. As can be seen from the graphs, the rate of rotation varies and is affected by transient presence of interactions of hemin side chains with G4 backbone and loops. The orange histogram in the left part indicates distribution of the phase dihedral angle for the relevant portion of trajectory. See Figure S6 for phase distribution summed over all simulations.

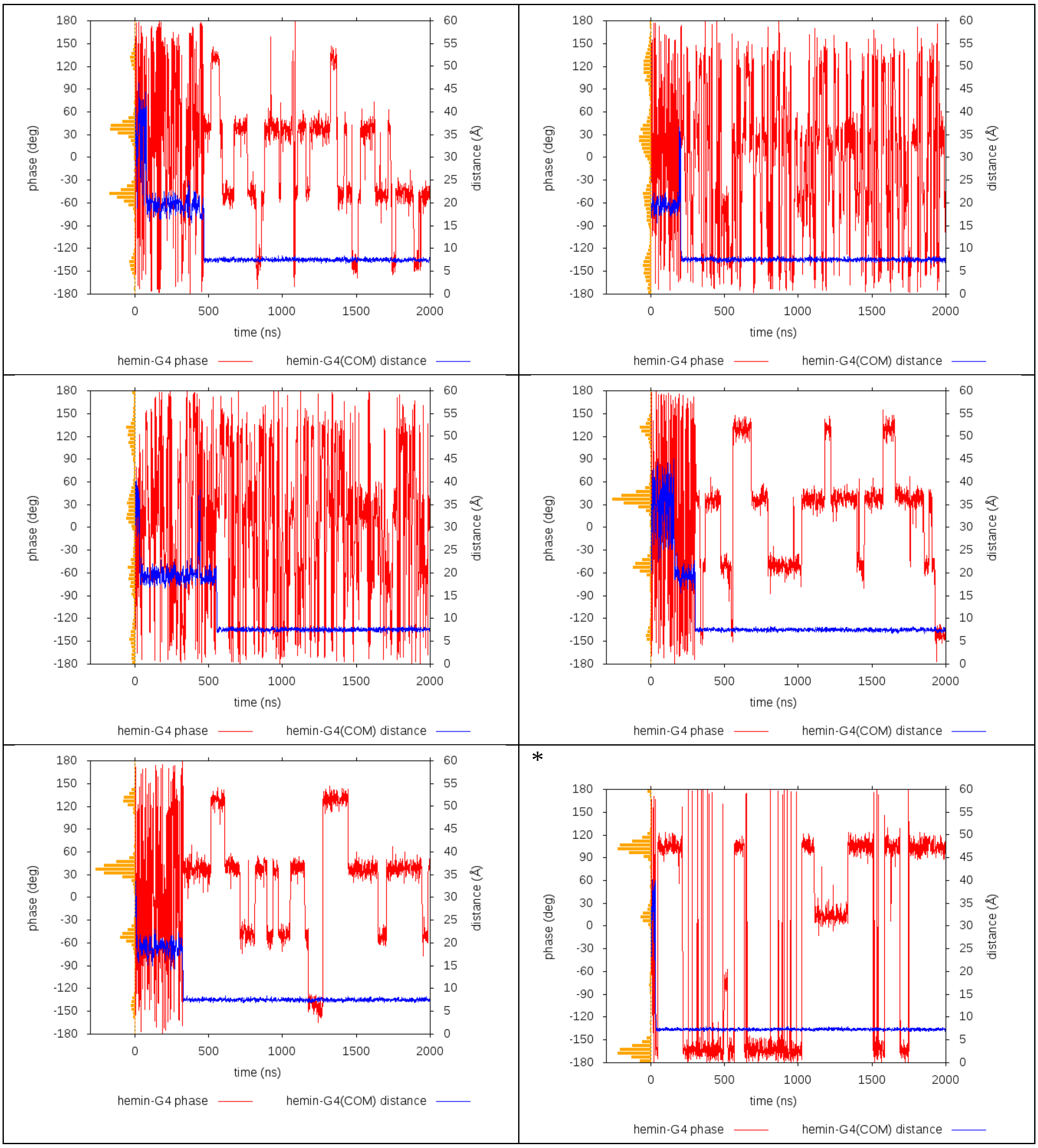

**Figure S5 (part 2/2)**. Hemin binding to 2LEE G4 and its rotation. See the first part for full explanation.

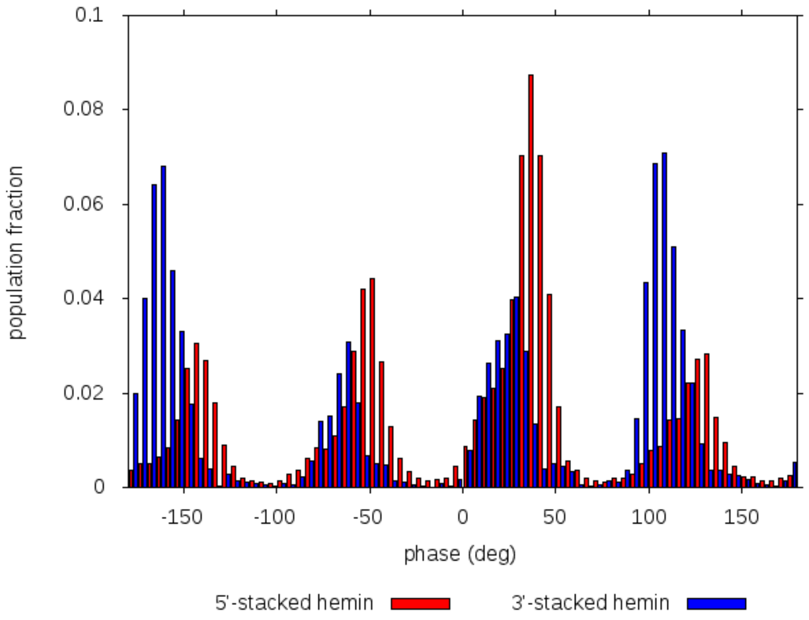

**Figure S6**. Rotational movement of hemin stacked on a quartet of 2LEE G4. Four-modal distribution was observed for hemin regardless of being bound to 5′-quartet or 3′-quartet. Bin width was set to 5 deg. The phase is an arbitrarily chosen dihedral angle as described in the legend to Figure S5. See the main text Figure 3C for 1KF1/hemin distribution. Figure S9 shows the distribution for the complex of tetramolecular G4 with hemin.

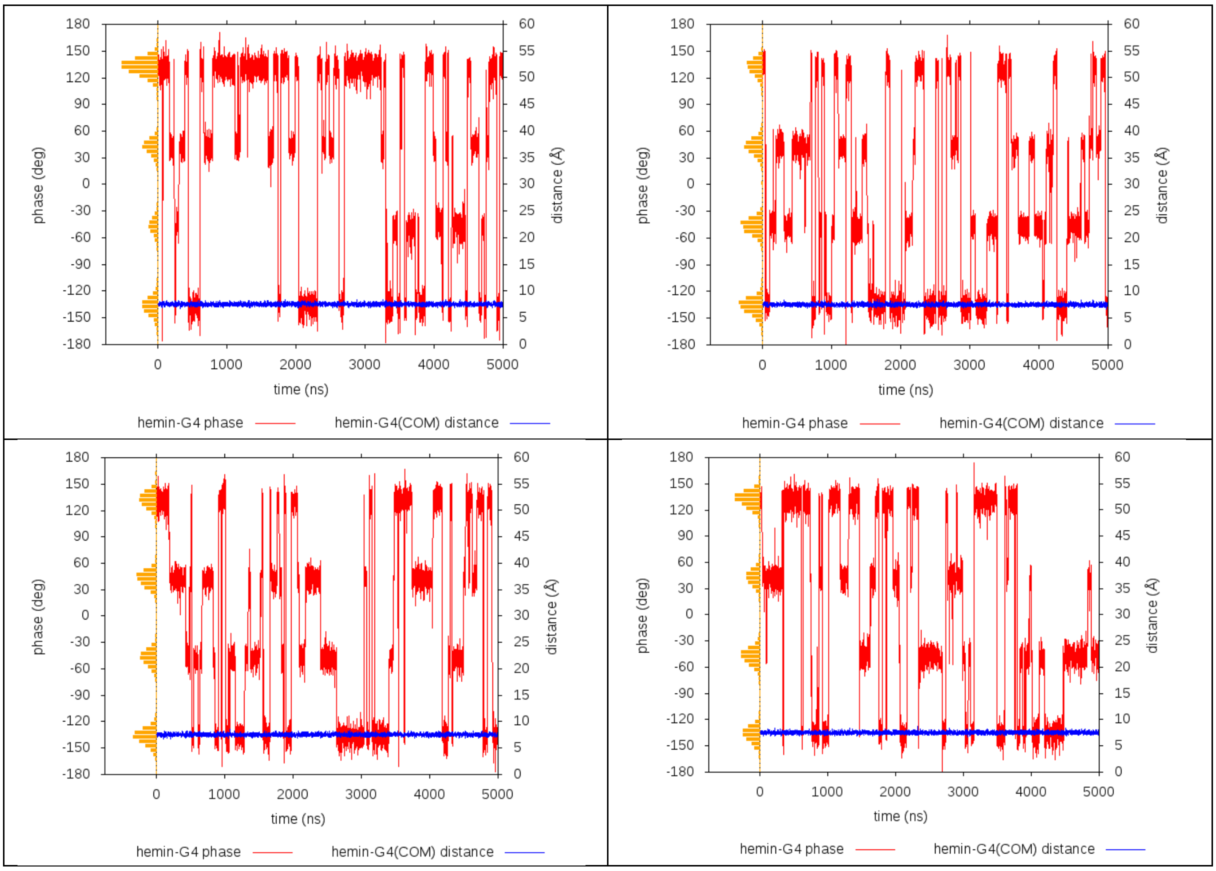

**Figure S7**. Hemin binding to the 5′-quartet of the tetramolecular G4 and its rotation in four simulations. Constant distance between hemin and G4 COM (blue line) indicates that hemin never unbound from the G4. The phase is an arbitrarily chosen dihedral angle defined by the following four points: 1) COM of C1′ atoms of G2 and G11 (i.e. Gs of the middle quartet from the first and last strand), 2) COM of base heavy atoms of guanines forming the middle quartet, 3) hemin Fe(III), 4) hemin ring hydrogen atom located between the two carboxyethyl side chains. It characterizes the rotation of hemin about the vertical axis. As can be seen from the graphs, the rate of rotation varies and is affected by transient presence of interactions of hemin side chains with G4 backbone. The orange histogram in the left part indicates distribution of the phase dihedral angle for the relevant portion of the trajectory. See Figure S9 for phase distribution summed over all simulations.

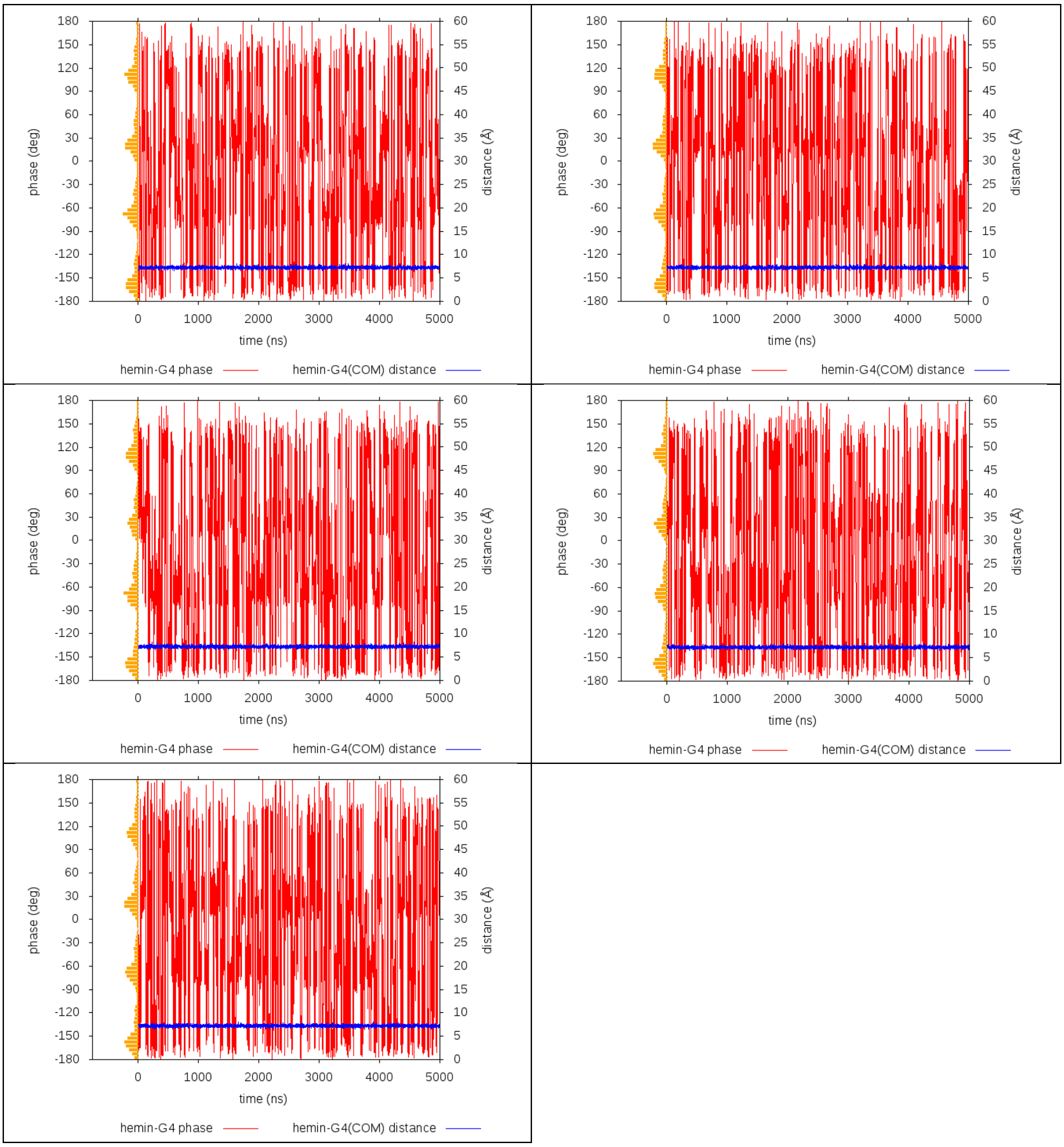

**Figure S8**. Hemin binding to the 3′-quartet of the tetramolecular G4 and its rotation in five simulations. Constant distance between hemin and G4 COM (blue line) indicates that hemin never unbound from the G4. The phase is an arbitrarily chosen dihedral angle defined by the following four points: 1) COM of C1′ atoms of G2 and G11 (i.e. Gs of the middle quartet from the first and last strand), 2) COM of base heavy atoms of guanines forming the middle quartet, 3) hemin Fe(III), 4) hemin side chain hydrogen atom located between the two carboxyethyl groups. It characterizes the rotation of hemin about the vertical axis. As can be seen from the graphs, the rate of rotation is significantly higher than in the complex with 5′-stacked hemin (Figure S7). The orange histogram in the left part indicates the distribution of the phase dihedral angle for the relevant portion of the trajectory. See Figure S9 for phase distribution summed over all simulations.

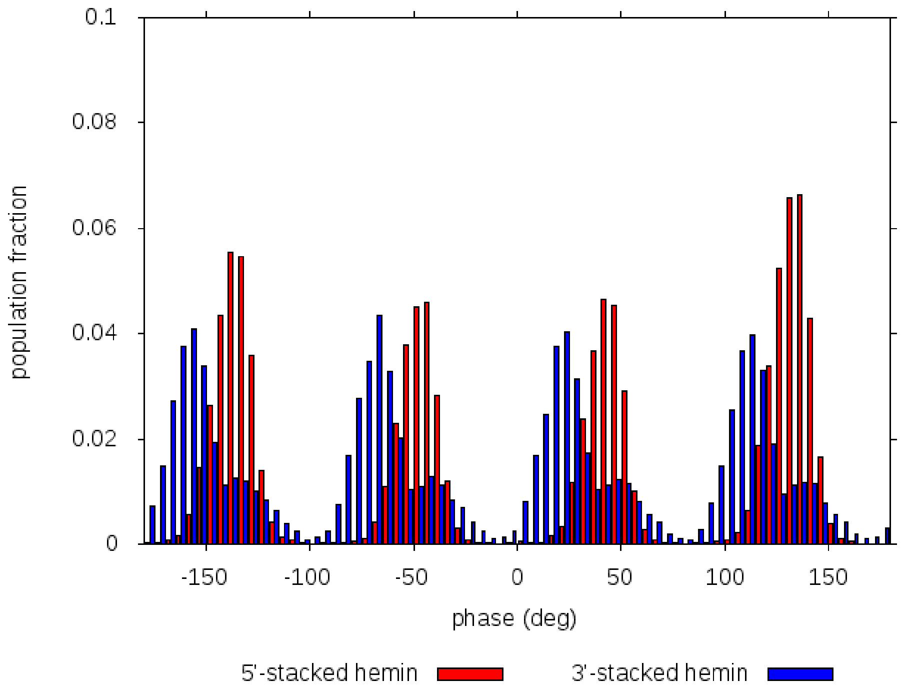

**Figure S9**. Rotational movement of hemin stacked on the 5′- and 3′-quartet of the tetramolecular G4. Four-modal distribution was observed for 5′-stacked hemin. The distribution has four peaks with four shoulders for 3′-stacked hemin. Bin width was set to 5 deg. The phase is an arbitrarily chosen dihedral angle as described in the legend to Figure S5. See the main text Figure 3C for 1KF1-hemin distribution and Figure S6 for the 2LEE-hemin distribution.

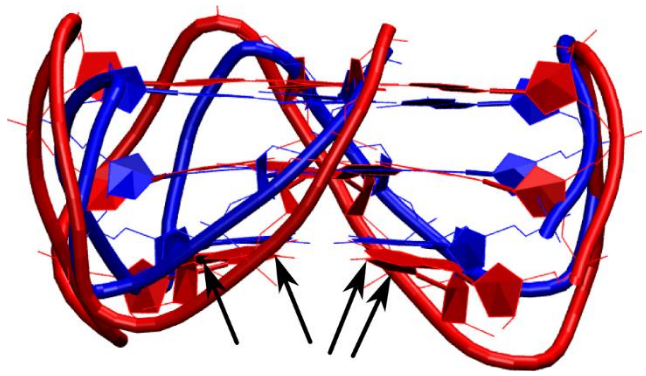

**Figure S10**. Hemin enforces quartet planarity when bound to the 3′-quartet. Average structure of the 3′-hemin/1KF1 complex is shown in blue (hemin not shown for clarity) and superimposed with the average structure of 1KF1 without hemin (red). The latter structure has virtually identical shape of the quartets as the average structure of 5′-hemin/1KF1 complex and the crystal structure. The black arrows indicate the four guanines in the 3′-quartet, which is visibly flattened upon hemin binding.

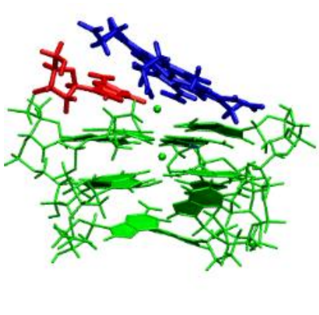

**Figure S11**. Guanine in the *t45* structure in vicinity of hemin iron. The iron in the center of hemin bound to the 5′-side of the tetramolecular G4 can be contacted by O6 atom of the top guanine. The pose is similar to 1KF1 with adenine-assisted transfer of hemin from the loop to the 5′-quartet (Figure 3B, middle structure). Hemin is blue, the involved guanine is red and G-stem with channel cations is green.

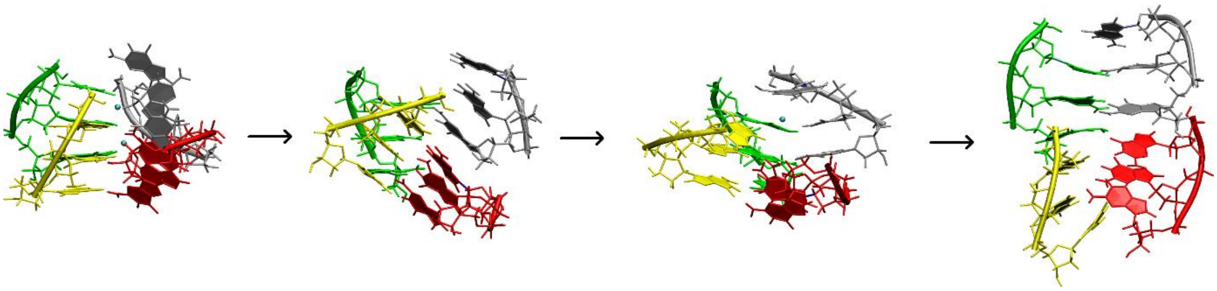

**Figure S12**. Unfolding intermediates of *t13-23* in the simulation without hemin. The first intermediate in this figure is where "further unfolding" is marked in Figure 5 in the main text. Each G-strand is shown in a different color. Channel cations (if present) are cyan. The unfolding process continued even further and the grey strand detached from the other three.

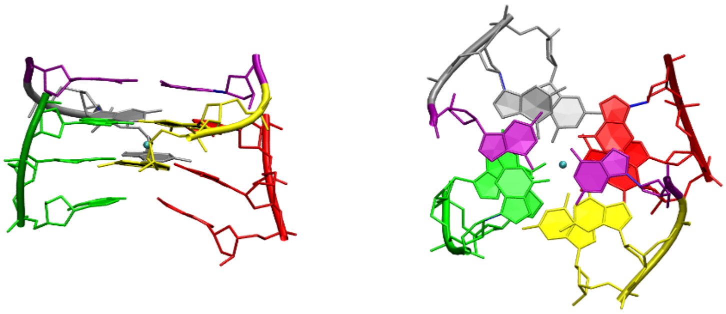

**Figure S13**. Slipped G4 *t25-45* stucture with a G:G base pair. Each G-strand is shown in a different color. Gs forming the *trans*-Watson-Crick base pair are shown in purple. Channel cation is cyan.

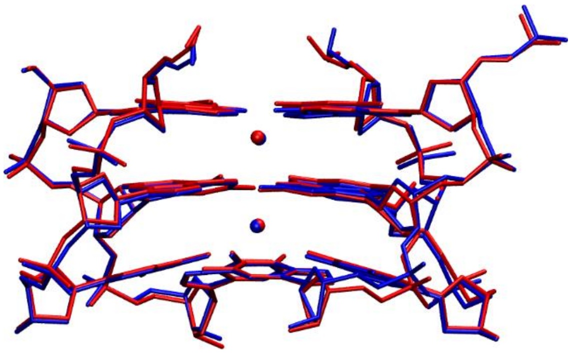

**Figure S14**. Iron(III) of hemin does not cause excessive inter-cation repulsion when bound. Average structure of G-stem with two channel $K^+$ at sites Q1 and Q2 in the large-box simulation *3a* (see Table S1) before hemin binding is red. Average structure of the same molecule after hemin binding to the 5′-quartet (top) is blue. Clearly, iron(III) of hemin does not exert any excessive repulsive force on the channel $K^+$ ions.
